# Comparative and systems analyses of *Leishmania* spp. non-coding RNAs through developmental stages

**DOI:** 10.1101/2021.05.17.444077

**Authors:** J. Eduardo Martinez-Hernandez, Victor Aliaga-Tobar, Carolina González, Rubens Monte-Neto, Alberto J. M. Martin, Vinicius Maracaja-Coutinho

## Abstract

*Leishmania* spp. is the etiological agent of leishmaniases, neglected diseases that seek to be eradicated in the coming years. The life cycle of these parasites involve different host and stress environments. In recent years, many studies have shown that several protein coding genes are directly involved with the development and host interactions, however, little is still known about the role of ncRNAs in life cycle progression. In this study, we aimed to identify the genomic structure and function of ncRNAs from *Leishmania* spp. and to get insights into the RNAome of this protozoan genus. We studied 26 strains corresponding to 16 different species of *Leishmania*. Our RNAome analysis revealed the presence of several ncRNAs that are shared through different species, allowing us to differentiate between subgenus as well as species that are canonically related to visceral leishmaniasis. We also studied co-expression relationships between coding genes and ncRNAs which in the amastigote developmental stage for *Leishmania braziliensis* and *L. donovani* revealed the presence of miRNA-like co-expressed with several coding genes involved in starvation, survival and histone modification. This work constitutes the first effort to characterize the *Leishmania* RNAome, supporting further approaches to better understand the role of ncRNAs in the gene regulation, infective process and host-parasite interaction.

## INTRODUCTION

Leishmaniases are a group of different diseases caused by *Leishmania* spp. parasites (1–3). These are classified based on their wide clinical manifestations, that range from self-cured skin lesions (cutaneous leishmaniasis or CL) to the visceral form (visceral leishmaniasis or VL) affecting organs such as liver, spleen and bone marrow (1,4). Reported annual infections vary from 0.7 to 1.3 million for CL and 300,000 cases for VL (5). Despite this high incidence and the numerous efforts to characterize these parasites molecularly, we still do not know how they control their gene expression.

The life cycle of *Leishmania* parasites involves invertebrate and mammal hosts, with several morphological changes and biochemical adaptations (6). Mammalian infection by *Leishmania* begins with the entry of metacyclic promastigotes (META) into the dermis by the bite of a sandfly, a blood-feeding Diptera part of the Phlebotominae clade. Metacyclic promastigotes are rapidly internalized by skin resident or recruited phagocytes notably neutrophils, macrophages and dendritic cells (7). Within the host cells, *Leishmania* parasites trigger the maturation and morphological transformation to amastigote (AMA) forms. At this point the parasites multiply by binary fission. This division leads to the lysis of the macrophage and induces the dissemination of amastigotes, allowing them to infect other cells (7–9). Finally, parasites re-enter into the vector by blood meal of an infected host, taking with them macrophages carrying amastigotes, triggering morphological changes to procyclic promastigotes (PRO) (10). Such a challenging life cycle requires parasite adaptation efforts to survive under nutritional starvation, sudden changes in temperature, pH and cellular plasticity varying from free living 15-30 μm length flagellated form to a 3-6 μm rounded intracellular form. All these can be achieved by a very well orchestrated post-transcriptionally controlled gene expression and epigenetic events as regulatory mechanisms already reported in trypanosomatids (11,12).

*Leishmania* parasites possess particular characteristics in the gene expression regulation (13). These features include a constitutive gene expression (14), polycistronic transcription (15–17) and post-transcriptional gene expression control (18). In addition, two recent studies in *L. donovani* reveal that gene dosis derived by aneuploidy modulates the gene expression (19,20). However, despite the fact that these works are mainly focused on characterizing regulatory mechanisms at the level of coding genes and have been described as participating in the development and pathogenesis of *Leishmania*, the role of other gene regulators such as non-coding RNAs has not been well studied, thus leaving a lack of understanding of how they participate in development regulation.

Non-coding RNAs (ncRNAs) are untranslated transcripts that participate modulating multiple biological processes, such as gene regulation at transcriptional (21) and post-transcriptional levels (22,23), developmental processes (24) or even in many diseases, such cancer (25) or pathogenic infections (26). ncRNAs can be classified into two main subclasses (25,27,28). The small RNAs (sncRNAs) (< 200 nt), a subclass that includes regulatory RNAs such microRNAs (miRNAs), small interfering RNAs (siRNAs), Piwi-interacting RNA (piRNAs), while the long non-coding RNAs (lncRNAs) (> 200 nt) includes several transcripts known to take place regulating several important processes in eukaryotes, such as genome imprinting (29), splicing (30) and chromatin organization (31). Many classes of regulatory ncRNAs have been virtually described in all eukaryotic cells since the miRNAs were firstly described in 1993 in *Caenorhabditis elegans* (32). Although ncRNAs have been extensively studied in recent years, little is known about their roles in the development and pathogenesis of eukaryotic microbes (33).

Since the first genome of *Leishmania major* was sequenced in 2005 (34), many studies addressed the understanding of the genome structure and function at protein coding genes level (34–43). In parallel, studies focused on ncRNAs in *Leishmania* parasites have increased since its first description in 2006 (44). These works have been mainly concentrated on the identification of specific ncRNAs classes, such as siRNA (45); microRNA-like and their regulatory roles (46); small RNAs derived from tRNAs and rRNAs as regulators of host-pathogen interaction processes (47); small nucleolar RNAs (snoRNA) and their function in rRNA processing (48); UTR-associated ncRNAs (uaRNAs) (49); or identifying lncRNAs and their putative functions (50). Recently, two works devoted to characterizing the ncRNAs repertoire in *L. braziliensis* brought new challenges on the understanding of non-coding transcripts’ role in trypanosomatids developmental life cycle stages (51,52). Certain lncRNAs act in *Trypanosoma brucei* differentiation and survival within the host (53). In this regard, a system biology approach has not yet been performed in *Leishmania* parasites.

In this work, we combined different computational, transcriptome, comparative genomics and systems biology approaches in order to identify and characterize the ncRNAs repertoire in 26 different strains belonging to 16 *Leishmania* species. We focused on the identification of regulatory ncRNAs that are shared or unique within different *Leishmania* species. We also analyzed the expression of these ncRNAs and their relationship with coding genes, in order to identify associated functions through coding/non-coding co-expression networks analysis. Interestingly, we observed several ncRNA-coding RNA pairs co-expressed in the same developmental stages. These pairs were differentially expressed (DE) in consecutive developmental steps, revealing the preponderant role of ncRNAs in the regulation of development and survival of *Leishmania* parasites.

## RESULTS

### Dissecting the repertoire of non-coding RNAs across *Leishmania* spp

We employed a combined computational approach based on covariance model comparisons of established RNA families, similarity searches using sequences retrieved from dozens of public databases and RNA-sequencing datasets to map the repertoire of ncRNAs available in 26 genomes from 16 *Leishmania* species. Our analysis characterized, for the first time, a plethora of different types of ncRNA in 16 species of *Leishmania*. The ncRNA predictions revealed a total of 274,737 ncRNAs, ranging from 5,149 (*Leishmania infantum* JPCM5) to 17,665 (*Leishmania turanica* LEM423). We identified 18 well known and 1 non-defined (unclassified) RNA classes spreaded over *Leishmania* spp. genomes, according to NRDR/NR2 (54) and Rfam v14 (55) original annotations and nomenclature; and an additional set of ncRNAs without any specific functional annotation but conserved according to other RNA sequences available in public databases and indexed in NR2. The most representative RNA class is miRNA, with an average of 1665 copies per genome. The less abundant class is scaRNA and ta-siRNA, presenting only 1 copy in LspLD974 and LtarParTarII, respectively. The top five RNA classes according to their average occurrence in all genomes were: 6,499 unclassified RNAs, 1,655 miRNAs, 1,051 sRNAs, 722 piRNAs and 399 rRNAs. miRNAs, sRNAs and unclassified RNAs correspond to around 87% of the ncRNAs found in each evaluated genome (Figure 1A). It is worth noting that the ncRNAs presented a total GC content ranging within 48% and 60%, and a length ranging from 18 (miRNAs) to 6,212 nt (unclassified RNAs). More information about the ncRNA repertory for each species is detailed in Supplementary Table 1 and Supplementary Table 2.

**Figure 1.**
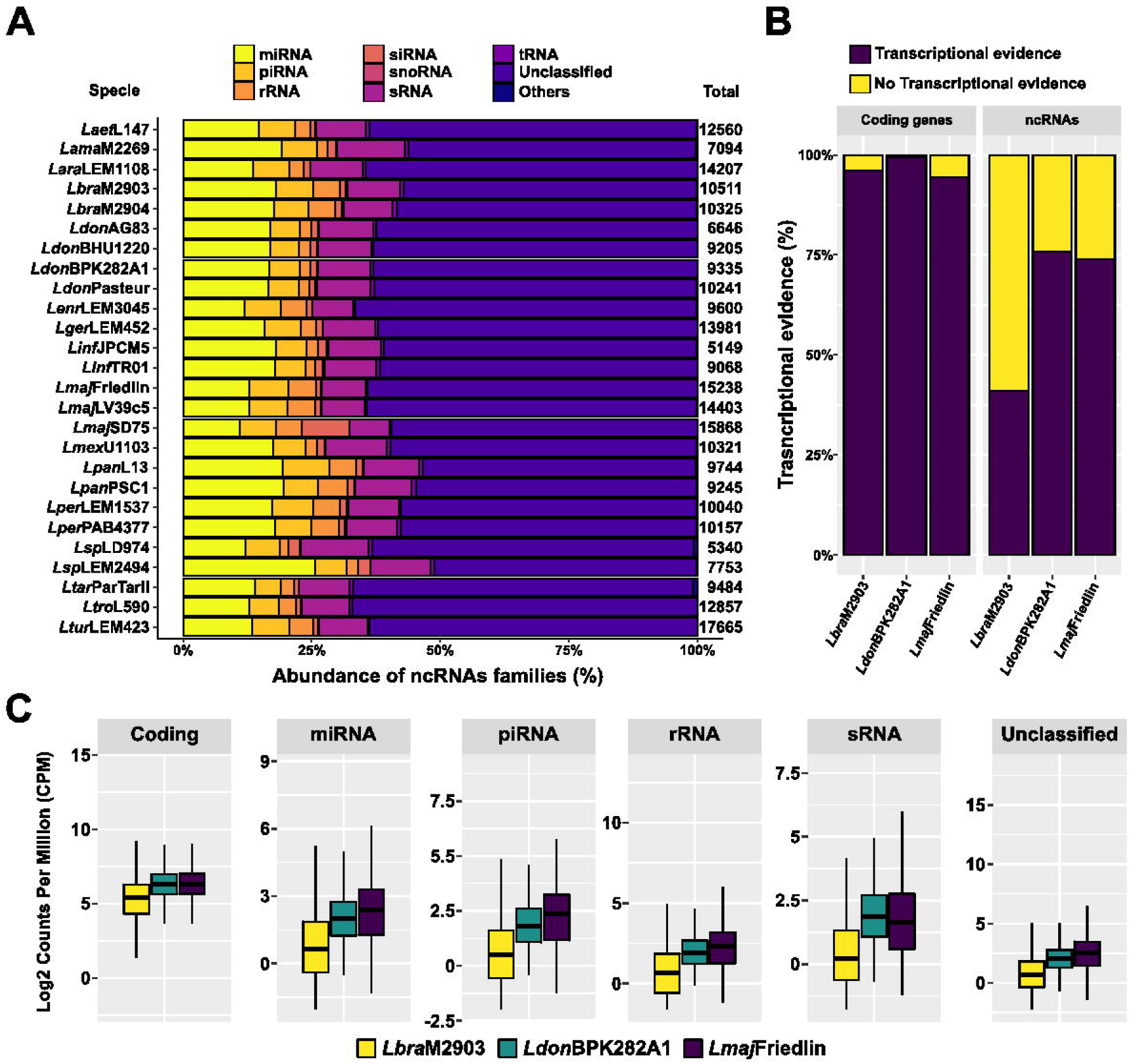
ncRNAs repertoire and transcriptional evidence in *Leishmania* parasites. **A**. Those ncRNAs who represent less than 0.5% were categorized as “Others”. The presence of a large proportion of non categorized ncRNAs, RNAs are elements that could not be assigned to any group due to lack of biological information related to their secondary structure or motifs. On the other hand, we also see the presence of miRNAs, piRNAs and sRNAS in large proportions. “Others”’ class represents all those ncRNAs that had less than 3% representation in the genomes of *Leishmania* spp. **B**. ncRNAs gene prediction was validated by transcriptional evidence using RNAseq analysis. We observed transcriptional evidence with a rate between 41.15% (*L. braziliensis*) to 75.89% (*L. donovani*). **C**. Comparative expression values of Log2 counts per million (log_2_CPM) normalized counts. Top five representative ncRNAs according to the number of predictions. We compare the expression of these subsets of ncRNAs against coding genes expressed in *L. braziliensis, L. donovani* and *L. major*; observing a clear trend of lower expression in ncRNAs versus coding genes.

To evaluate the transcriptional evidence for predicted ncRNAs, we integrated publicly available RNA-seq data of three strains from three different species: *L. braziliensis* M2903 (MHOM/BR/75/M2903), study accession PRJNA494068 published by Ruy and cols. (52); *L. donovani* BPK282A1 (MHOM/NP/02/BPK282), study accession PRJEB15610 published by Dumentz and cols. (19) and *L. major* Friedlin (MHOM/IL/81/Friedlin), study accession PRJNA252769 published by Fernandes and cols. (56). After applying an expression cut-off of 0.5 counts per million (CPM) to be considered as a *bona fide* expressed transcript, we were able to obtain transcriptional evidence for a number of ncRNAs that range between 41.15% for *L. braziliensis* M2903 to 75.89% of the predicted ncRNAs for *L. donovani* BPK 282. In contrast, we validated the expression of around 97% of protein coding genes in these three species using the same RNA-seq datasets (Figure 1B).

After filtering all normalized values of the top representative ncRNAs, we compared expression values measured as log_2_ CPM of both all ncRNAs and all protein coding genes for each strain (Figure 1C). We identified a mean value of CPM for ncRNAs ranging from 2.55 for piRNAs to a maximum of 84.3 for unclassified RNAs in *L. braziliensis*. We also observed a mean CPM value of 70.07 for coding genes. The trend is similar for ncRNA expression in *L. donovani*, with an average expression that ranges between 6.50 for sRNA to 103.61 for rRNAs using log_2_ CPM values, while coding genes showed an average expression value of 118.32 CPM. In the same way, we noticed that the behavior of both coding genes and ncRNAs in *L. major* is similar to the other two previously mentioned species, having a value that ranges from 6.32 for sRNAs to 10.05 for unclassified RNAs; while coding genes displayed an average expression value of 107.97 log CPM.

### The *Leishmania* spp. pan-RNAome: ncRNAs conservation across 26 genomes

We carried out a comparative genomic analysis to find patterns of conservation throughout the ncRNA repertoire in all *Leishmania* genomes. After a clusterization process using CD-HIT (57) the primary sequences of all identified ncRNAs using an inventory of 274,737 ncRNAs, were distinguished into 5,482 clusters, named the pan-RNAome of Leishmania spp. from now on. The pan-RNAome was divided into a core-RNAome, formed by 436 (7.95%) clusters (Figure 2A), representing 196,618 ncRNAs shared among all species, and a total of 5,046 (92.05%) clusters of 75,878 dispensable ncRNAs or accessory-RNAome, shared within 1 to 25 genomes (Figure 2A). From the accessory-RNAome, we identified a set of 2,241 ncRNAs (1,962 clusters) unique for each particular evaluated *Leishmania* genome, corresponding to 35.79% of the pan-RNAome (Figure 2A).

**Figure 2.**
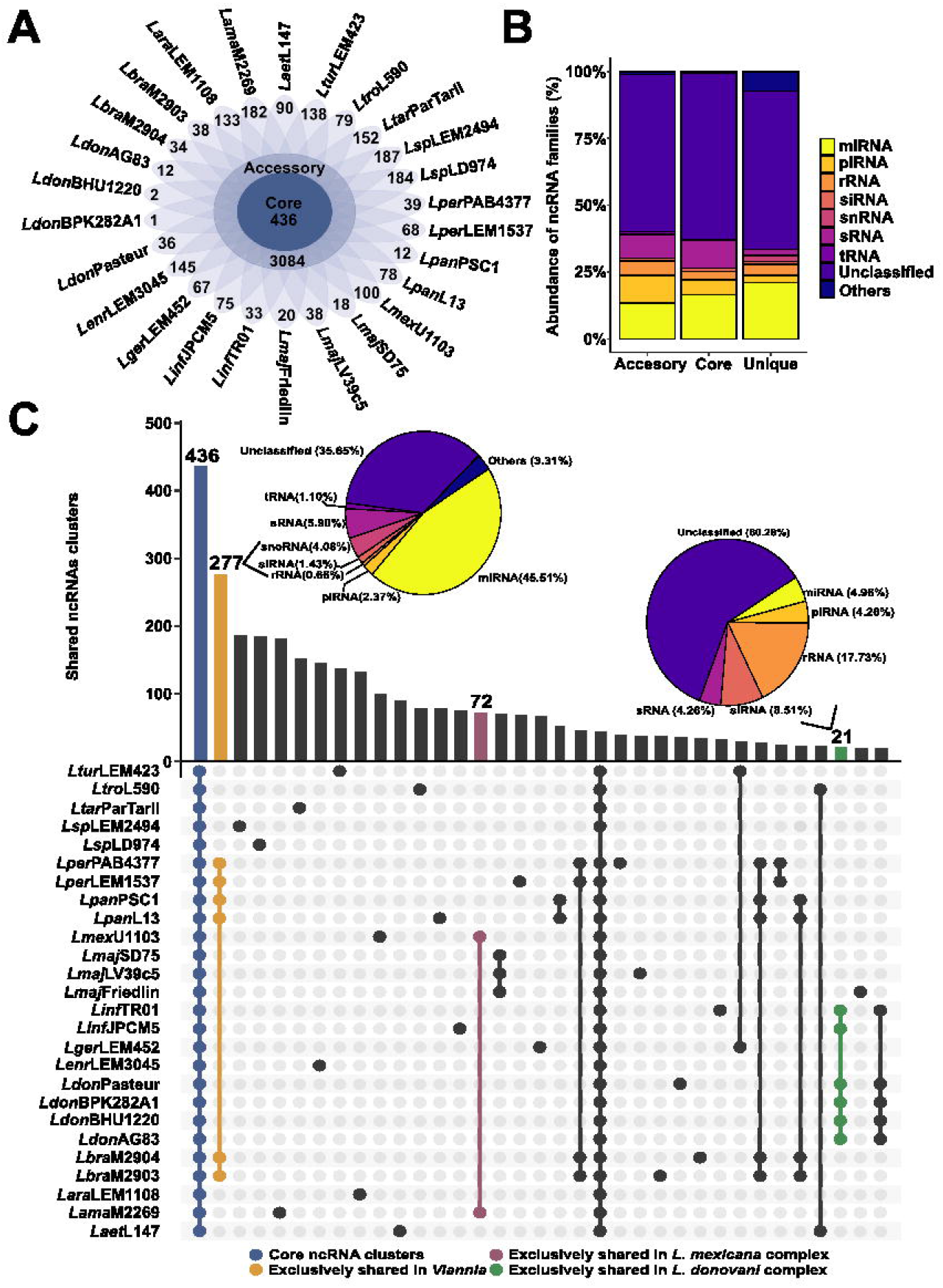
Conservation analysis through *Leishmania* spp. **A**. Flower map indicates the conservation of all ncRNA in *Leishmania* parasites. Central circle represents the core RNAome. Accessory ncRNA is represented in the second circle and each petal contains unique ncRNA found for each evaluated strain. **B**. Abundance of ncRNAs families in pan-RNAome of *Leishmania* spp. **C**. Upset plot corresponds to the top 35 interactions between ncRNAs in all 26 species. Interactions in green represent all clusters conserved exclusively in species related to visceral leishmaniasis (VL) (*L. donovani* and *L. infantum* strains) related species. Unique interactions in the *Viannia* subgenus (*L. braziliensis, L. panamensis* and *L. peruviana* strains) are represented in orange. *Leishmania mexicana* complex (*L. mexicana* and *L. amazonensis*) clusters are represented in purple. Core interactions are represented in blue dots. Pie charts represent the abundance of ncRNAs classes in *Viannia* and VL related species.

Regarding ncRNA types, miRNAs, sRNAs, piRNAs, rRNAs, siRNAs, tRNAs and snoRNAs were the major classes that compose the 37,48% of the core-RNAome, the remaining 62,52% corresponds to unclassified RNAs (Figure 2B). The subset of unique ncRNAs is the most diverse in terms of ncRNA types. Nonetheless, the most common classes were unclassified RNAs, miRNAs, rRNAs, gRNAs and piRNAs, which together represent 90,59% of the total inventory for this subset (Figure 2B). Notably, we also identified a subset of accessory ncRNAs that allowed us to distinguish between *Viannia* and *Leishmania* subgenus. Species belonging to the *Viannia* subgenus shared 277 clusters of ncRNAs absent in the other groups (Figure 2C). These *Viannia* subgenus-specific clusters are mainly annotated as gene expression regulatory ncRNAs, with 56% part of either lncRNAs, miRNAs, piRNAs, sRNAs and siRNAs (Figure 2C). Next, we evaluated the ncRNAs conservation in species canonically related to VL. For that, based on the literature, we selected *Leishmania donovani* and *Leishmania infantum* as representative of VL classically associated species (2,58,59), where we found a subset of 21 ncRNA clusters (Figure 2C), which belongs to unclassified RNAs, rRNAs, siRNA, miRNAs, piRNAs and sRNAs (Figure 2C).

To evaluate the relationship between the phylogeny based on protein coding genes and ncRNA classes distribution within all *Leishmania* species that cause different clinical manifestations or subgenus, we compared the phylogenetic relationship for the primary sequences of 613 core orthologous genes conserved within all 26 *Leishmania* spp. isolates, with a clustering generated with Jaccard similarity computed using the ncRNA presence/absence matrix. Interestingly, both approaches based on sequence phylogeny and Jaccard coefficient clustering, showed a similar behavior, demonstrating a phylogenetic relation in the conservation and distribution of ncRNAs (Supplementary Figure 1).

### Identification of RNA interference pathway in *Leishmania* genomes

To provide potential evidence for the presence of RNAi machinery in *Leishmania* genomes (*L. braziliensis, L. donovani* and *L. major*), we applied a computational strategy based on functional domain preservation and remote sequence homology. For this, we surveyed Pfam IDs for the canonical and non-canonical miRNA/RNAi pathways and performed a domain search with their respective hidden Markov models (HMMs). This approach allowed us to identify Argonaute-like (AGO-like) proteins in *L. braziliensis, L. donovani* and *L. major* (Table 1). Also, with this search scheme it was possible to identify a Dicer-like protein in *L. braziliensis*, but not in *L. donovani* or *L. major*. Likewise, Exportin 5, RAN (GTP-binding nuclear protein Ran) and Exportin 1 were identified in all three genomes. To determine the presence of AGO-like proteins in *L. donovani* and *L. major*, we performed a BLAST alignment of all AGO-like proteins reported in TritrypDB against both genomes. The search yielded the presence of a locus that corresponds to 342 nt (114 aa) in chromosome 11 of *L. donovani* HU3 strain, that match with LDHU3_11.0760 protein that is annotated as hypothetical, but is related to AGO-like proteins with more than 50% of identity, 77% of query coverage and e-values <1e-10 in our BLAST outcomes, however in L. donovani BPK282A1, we identify a pseudogene corresponding to AGO-like protein. We identified an AGO-like protein in *L. major* (LmjF.11.0570) with an identity of more than 50% and a query coverage > 98%. Here, we observed only a small portion of AGO1 protein described as pseudogene in *L. donovani* and *L. major* species and a functional form in *L. braziliensis* (Figure 3A). Unfortunately, Drosha, DGCR8 and TRBP proteins of the canonical RNAi pathway were not found in any of the three genomes evaluated, which leads us to think that a non-canonical pathway is present in *Leishmania* parasites. Additional analysis should be carried out in order to determine most effectively the possible pathway of siRNA/miRNA biogenesis in these species. Table 1 lists the proteins that act in the canonical and non-canonical RNAi pathway in *Leishmania* likewise, the expression patterns for elements of both pathways are described in Figure 3B.

**Table 1.** RNAi components in different *Leishmania* species.

| Protein | Species |  |  |
| --- | --- | --- | --- |
|  | *L. braziliensis* | *L. donovani* | *L. major* |
| Canonical pathway |  |  |  |
| AGO1 | LBRM2903_110008400.1 | LDHU3_11.0760**<br>LdBPK_110500.1* | LmjF.11.0570* |
| DGCR8 | - | - | - |
| Dicer | LBRM2903_230009800.1 | - | - |
| Drosha | - | - | - |
| Exportin 5 | LBRM2903_350037400.1 | LdBPK_362810.1 | LmjF.36.2680 |
| TRBP | - | - | - |
| RAN | LBRM2903_250023400.1 | LdBPK_251460.1 | LmjF.25.1420 |
| Non-canonical pathway |  |  |  |
| Exportin 1 | LBRM2903_320017800.1 | LdBPK_321160.1 | LmjF.32.1100 |
| tRNAse Z | LBRM2903_220016000.1 | LdBPK_220940.1 | LmjF.22.1120<br>LmjF.22.1130 |
| METTL1 | LBRM2903_340062500.1 | LdBPK_355200.1 | LmjF.35.5230 |

pseudogene = \* gene of *L. donovani* H3=\*\*

**Figure 3.**
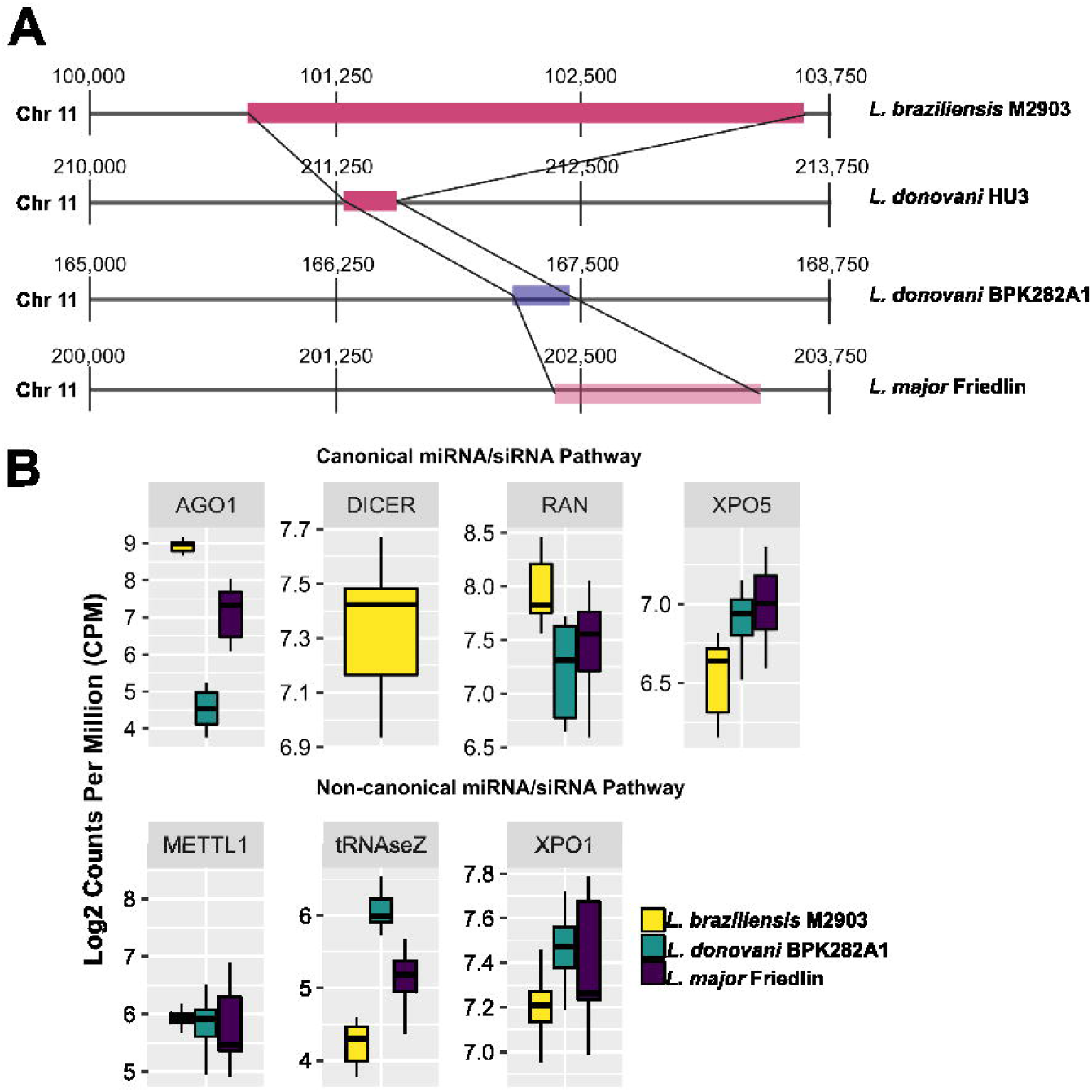
Synteny of AGO1-like gene in Leishmania species and expression patterns of canonical and non-canonical miRNA/siRNA pathway. **A**. AGO1-like genes are present in chromosome 11 in *Leishmania* species. In *L*.*braziliensis* M2903 and *L. donovani* HU3 are a protein coding gene (solid pink), however in *L. donovani* BPK282A1 and *L. major* Friedlin (transparent pink) were detected as a transcribed pseudogene. In addition, we observed that this pseudogene in *L*.*donovani* BPK282A1 is in a 3’ to 5’ arrangement (transparent blue). **B**. Expression patterns of the genes involved in canonical and non-canonical miRNA/siRNA pathway.

### Differential expression of coding genes and ncRNAs

In order to retrieve further information of the potential participation of these ncRNAs in key biological processes, we obtained the set of differentially expressed coding genes and ncRNAs between the three main developmental stages of *L. braziliensis, L. donovani* and *L. major*. We analyzed the expression of 4,325 ncRNAs and 8,908 coding genes of *L. braziliensis*, in pairwise comparison through amastigote, metacyclic promastigote and procyclic promastigote developmental stages(Figure 4A). After this analysis, we identified a total of 1,103 differentially expressed (DE) genes overexpressed at the amastigote stage, with 783 mRNAs and 220 ncRNAs exclusively overexpressed in amastigotes. Another 537 genes (466 mRNAs, 71 ncRNAs) were shared with metacyclic, meanwhile 26 (6 mRNAs, 20 ncRNAs) were shared with the procyclic stage. In contrast, for metacyclic we obtained 117 overexpressed ncRNAs, and 212 protein coding genes. For the procyclic developmental stage, we detected the overexpression of 503 coding genes and 378 ncRNAs. Additionally, we found 316 (179 mRNAs, 137 ncRNAs) shared DE genes within metacyclic and procyclic stages (Figure 4B, Supplementary File 1). In a similar way, we evaluated the differential expression in *L. donovani* of 7,085 ncRNAs and 7,998 coding genes. Compared gene expression patterns between the amastigote and promastigote developmental stages identified 777 genes (382 ncRNAs, 395 coding genes) overexpressed in promastigote, and 623 (78 ncRNAs, 545 coding genes) overexpressed amastigote (Supplementary Figure 2A, Supplementary File 1). Finally, we determined the differential expression of ncRNAs and coding genes from *L. major*, identifying 331 DE genes (222 ncRNAs, 109 coding genes) overexpressed exclusively in the metacyclic stage, and 219 (96 ncRNAs, 123 coding genes) overexpressed only in the procyclic stage (Supplementary Figure 3A, Supplementary File 1). To reveal the biological processes involved in development of *Leishmania* parasites,

**Figure 4.**
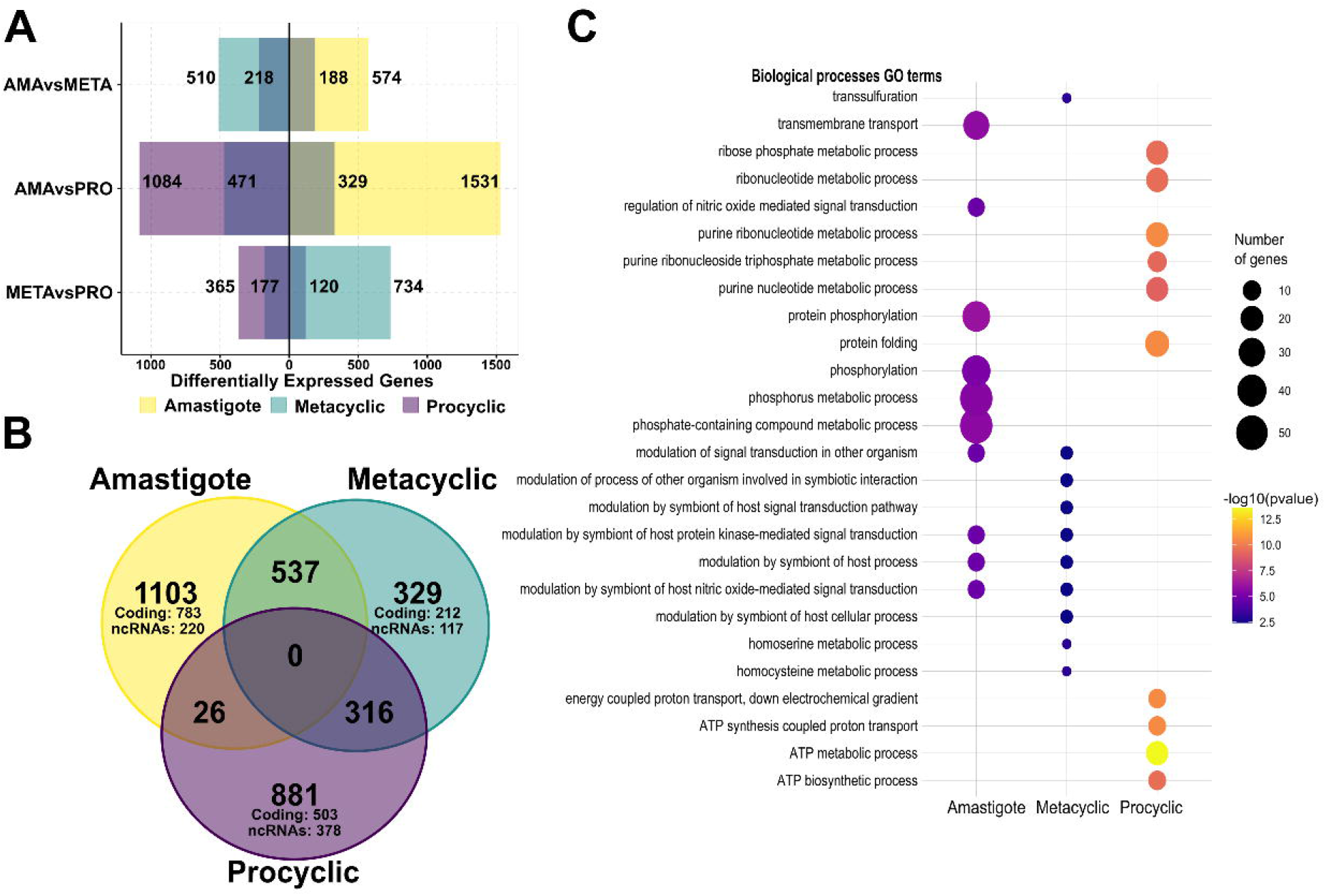
Differentially expressed coding and non-coding genes through developmental stages in *L. braziliensis*. **A**. Pairwise comparison of gene expression in Amastigote, Metacyclic and Procyclic developmental stages. The number on top of each bar represents the number of all overexpressed genes in each stage comparison. The shadow bar represents the ncRNAs overexpressed. **B**. Venn diagram to identify exclusively DEGs of each developmental stage. **C**. These exclusive genes were used to identify the enrichment of GO biological processes. Here are the top 10 categories according to p-value for each developmental stage.

we then studied the functional annotation of the DE mRNAs. We considered functional terms to be enriched if they sowed a *p* value < 0.001 in Gene Ontology analysis using TritrypDB annotation tools. DE mRNAs in *L. braziliensis* amastigotes were significantly enriched in GO molecular function terms related to protein phosphorylation (GO:0006468), transmembrane transport (GO:0055085), modulation by symbiont of host process (GO:0044003), modulation of signal transduction in other organism (GO:0044501) or regulation of nitric oxide mediated signal transduction (GO:0010749) (Figure 4C, Supplementary File 2). Meanwhile, metacyclic biological processes showed that DE mRNAs were enriched in terms related to metabolic process of homocysteine (GO:0050667) and homoserine (GO:0009092), transsulfuration (GO:0019346), modulation by symbiont of host protein kinase-mediated signal transduction (GO:0075130) or modulation by symbiont of host nitric oxide-mediated signal transduction (GO:0044081). In contrast, procyclic biological processes exhibited an enrichment in energy metabolism related terms, such ATP metabolic process (GO:0046034) or ATP synthesis coupled proton transport (GO:0015986). Additionally, categories involved in ribonucleotide metabolism, such as purine ribonucleotide metabolism (GO:0009150), ribose phosphate metabolism (GO:0019693) or ribonucleotide metabolism (GO:0009259) were enriched in *L. braziliensis* promastigotes (Figure 3B, Supplementary File2). The enrichment analysis of biological processes involved in development of amastigote and promastigote in *L. donovani*, showed DE mRNAs related to cellular nitrogen compound metabolic process (GO:0034641), peptide biosynthetic process (GO:0043043), ncRNA processing (GO:0034470) or ribosome biogenesis (GO:0042254) significantly enriched in amastigotes. Promastigote DE mRNAs in *L. donovani* presented significant enrichment in processes related to energy metabolism, such as pyruvate metabolism (GO:0006090), glycolytic process (GO:0006096) or ATP metabolism (GO:0046034) (Supplementary Figure 2B, Supplementary File 2). Furthermore, metacyclic promastigotes in *L. major* showed a biological processes enrichment in terms linked to signal transduction (GO:0007165) and cellular response to stimulus (GO:0051716), while procyclic promastigotes in *L. major* present, similarly to the procyclic promastigotes in *L. braziliensis*, an enrichment in terms related to energy metabolism (ATP biosynthetic process (GO:0006754), energy coupled proton transport, down electrochemical gradient (GO:0015985)) and nucleotide metabolism (purine nucleoside triphosphate biosynthetic process (GO:0009145); ribonucleoside triphosphate metabolic process (GO:0009199)). Additional information and the complete list of enriched terms were made available in Supplementary Figure 3B and Supplementary File 2.

### Co-expression network analysis identifies developmental stages-associated gene modules in *Leishmania* spp

A total of 13,233 genes (sum of coding and ncRNAs) of *L*.*braziliensis*, 15,082 of *L. donovani*, and 19,406 genes of *L. major* were employed to construct the weighted gene co-expression networks using WGCNA (60). The set up condition by our network’s construction was a soft threshold power β at 14 (for *L. braziliensis* and *L. donovani*) and 22 (for *L. major*), the scale-free network fitting index (R^2^) greater than 0.85 was setted to ensure low mean connectivity and high scale independence. Seventeen co-expression modules were recognized in *L. major*, in contrast with the eighteen present in *L. donovani* and twenty two in *L. braziliensis*. The number of genes per module varies from 55 (M19) to 2,539 (M3) in *L. braziliensis*. In *L. donovani*, it ranges from 1 (M0) to 4,896 (M3) genes; and in *L. major* we found between 74 (M10) to 5,641 (M12) genes per module. The total number of genes per module, as well as its composition of ncRNAs and coding is described in detail in Supplementary Table 3.

To analyze the correlation of each module in different *Leishmania* spp. developmental stages, we used a module-development stage relationship comparison. The relationship between co-expression modules and developmental stages is shown in Figure 5 and Supplementary Figure 4. After assessing strong correlations between all modules and developmental stages in *Leishmania* parasites, we found that module M2 had the highest correlation with the procyclic promastigote stage in *L. braziliensis* (R^2^ = 0.87 and *p* < 0.001). In the same way, the module M6 has stronger correlation with metacyclic promastigote (R^2^ = 0.9 and *p* < 0.001), and M18 with amastigote (R^2^ = 0.96 and *p* < 0.001) (Figure 5). For *L. donovani*, we observed that the highest module associated with the promastigote developmental stage was M11 (R^2^ = 0.95 and P < 0.001), and M5 to amastigote (R^2^ = 0.95 and *p* < 0.001) (Supplementary Figure 4A). In addition, modules M2 (R^2^ = 0.81 and *p* < 0.001) and M9 (R^2^ = 0.65 and *p* < 0.001) presented the highest correlation value for procyclic promastigote and metacyclic promastigote developmental stages, respectively in *L. major* (Supplementary Figure 4B).

**Figure 5.**
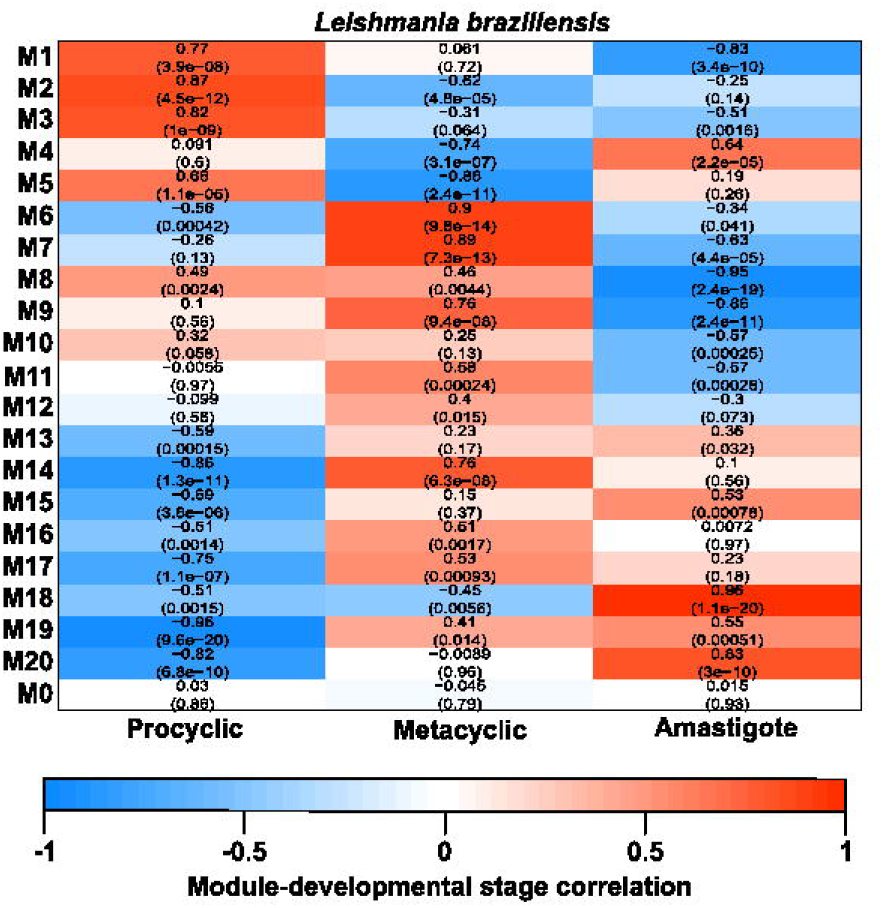
Co-expression module-Developmental stages associations. Each row corresponds to a module, column to a developmental stage. Each cell contains the corresponding correlation and p-value (parenthesis). The table is color-coded by correlation according to the color legend. **A**. *L. braziliensis*. Here we identify that modules M1 (R^2^= 0.77 and *p* < 0.001), M2 (R^2^= 0.87 and *p* < 0.001) and M3 (R^2^= 0.82 and *p* < 0.001) have the major correlation with the procyclic developmental stage, besides M6 (R^2^= 0.90 and *p* < 0.001), M7 (R^2^= 0.89 and *p* < 0.001), M9 (R^2^= 0.76 and *p* < 0.001) and M14 (R^2^= 0.76 and *p* < 0.001) are tightly related to the metacyclic stage and modules M18 (R^2^= 0.96 and *p* < 0.001) and M20 (R^2^= 0.83 and P < 0.001) are correlated to the amastigote developmental stage.

### Detection of potential key miRNA-like involved in amastigote life cycle in *Leishmania* parasites

In order to assess the possible functions of the identified miRNA-like in amastigote developmental stage, DE mRNAs and miRNAs-like were selected to filter the co-expression network corresponding to this life cycle stage in *L. braziliensis* and *L. donovani*. Next, we predicted the functions of selected miRNAs-like from the coexpression network by combining hub- and module-based methods previously reported (61). Here, we selected the top ten hubs presenting miRNA-like RNAs based on their centrality degree, expression and gene significance metric. According to this approach, a final set of top ten miRNAs-like presented co-expression with 89 DE mRNAs in *L. braziliensis* amastigotes (Figure 6A). At the same time, the evaluation of the top ten miRNA-like in *L. donovani* amastigotes showed that these co-expressed with 206 protein coding genes (expressed exclusively in amastigote developmental stage - Figure 6B). Each cluster presenting the above mentioned DE mRNAs were used to conduct GO enrichment analysis in order to try to obtain potential associations of these miRNA-like RNAs with known biological processes to these top in each species. In Figure 6C, we included the top 10 enriched terms for each cluster for both species (Supplementary File 3). The main GO biological processes terms enriched, according to their *p values*, in *L. braziliensis* were related to molecule transport (transmembrane transport (GO:0055085), transport (GO:0006810, sulfur compound transport (GO:0072348)), inorganic anion transport (GO:0015698)) or response to nutrient levels (cellular response to starvation (GO:0009267), response to starvation (GO:0042594), response to nutrient levels (GO:0031667)) (Figure 6C and Supplementary File 3). Meanwhile, in *L. donovani* we observed that the main GO terms enriched were related to amide and peptide metabolism biosynthetic processes (peptide biosynthetic process (GO:0043043), amide biosynthetic process (GO:0043604)), as well gene expression (GO:0010467) and translation (GO:0006412) (Figure 6C and Supplementary File 3).

**Figure 6.**
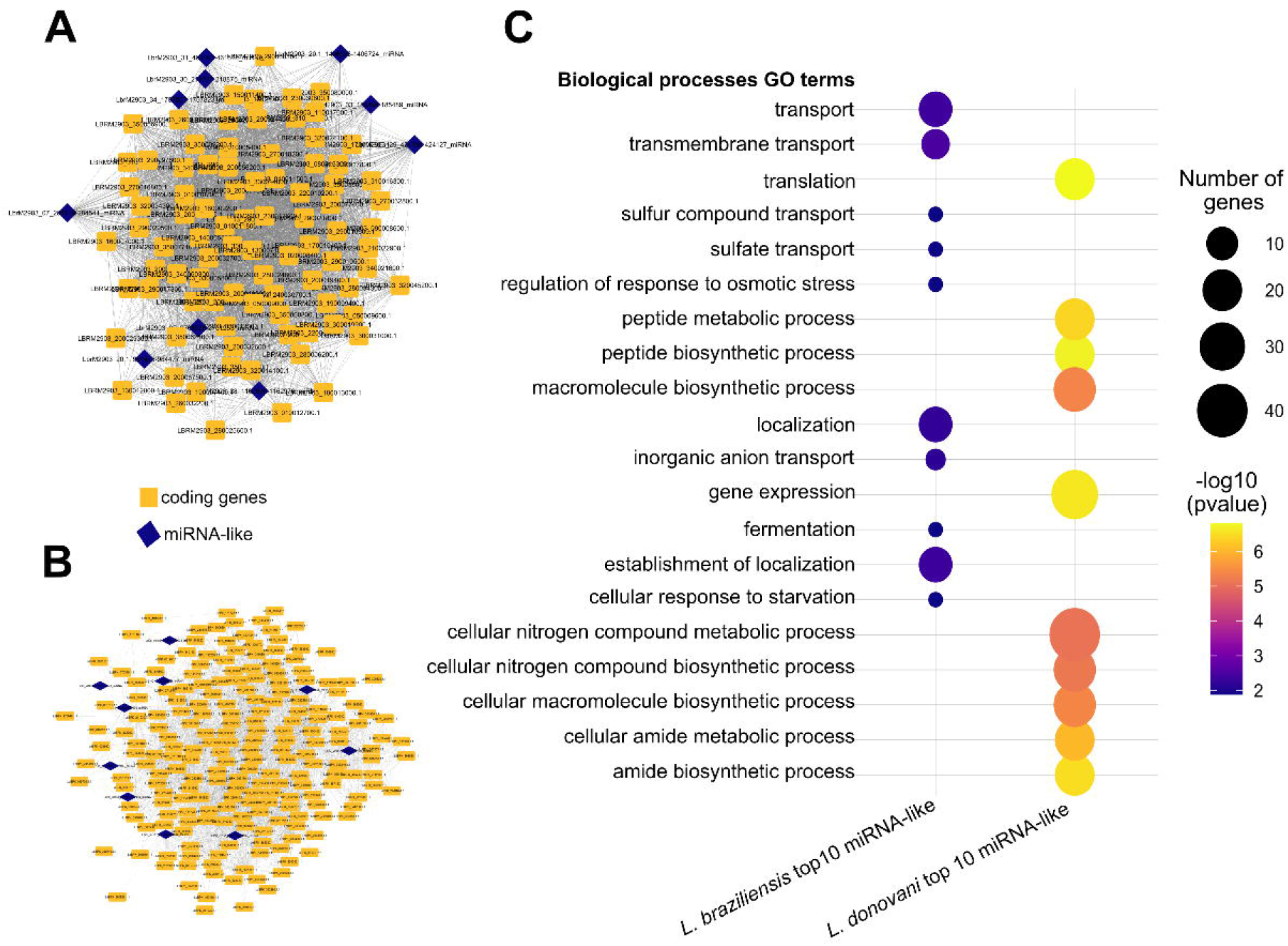
Biological processes Go enrichment of top 10 miRNA-like related to amastigote developmental stage in *L. braziliensis* and *L. donovani*. **A** and **B**. Sub-network of top 10 miRNA-like mainly related amastigotes in *L. braziliensis* (**A**) and *L. donovani* (**B**). The blue triangles represent miRNA-like genes. The orange squares represent the protein-coding genes that are co-expressed with it. **C**. Top 10 biological processes terms for functions assigned to the miRNAs-like in each species.

Through the guilty by association functional annotation approach, we predicted the possible function of the top miRNA-like that participates in amastigote developmental stage in *L. braziliensis* and *L. donovani*. The LbrM2903_20.1_984445-984477_miRNA miRNA-like gene was co-expressed with 40 coding genes (Figure 4A). Among them, one protein is annotated as target for rapamycin kinase (TOR) kinase 3 (TOR3), encoded by LBRM2903_200049400.1. This protein was related mainly to nutrient starvation (Supplementary File 3). We predicted that this miRNA-like would participate in biological processes mainly associated with response to nutrient levels, cellular response to external stimulus and regulation of response to stress, *p*-values<0.05, (Figure 7D and Supplementary File 3). We also observed that Ld17_v01s1_485484-485505_miRNA was co-expressed with 86 coding genes (Figure 7B), and their predicted biological processes activity was associated with gene expression, ncRNA metabolic process, and DNA synthesis involved in DNA repair (Figure 6D and Supplementary File 3). Additionally, we identified that Ld27_v01s1_883438-883470_miRNA co-expressed with other 17 protein coding genes in *L. donovani* amastigotes (Figures 7C) possessing an enrichment in histone deacetylation process associated with the gene LdBPK_241410.1, that encodes to a NAD-dependent histone deacetylase. Interestingly, this miRNA-like was co-expressed with Histone H2B and PriPol-like-2, a protein involved in DNA replication during stress conditions (Supplementary File 3).

**Figure 7.**
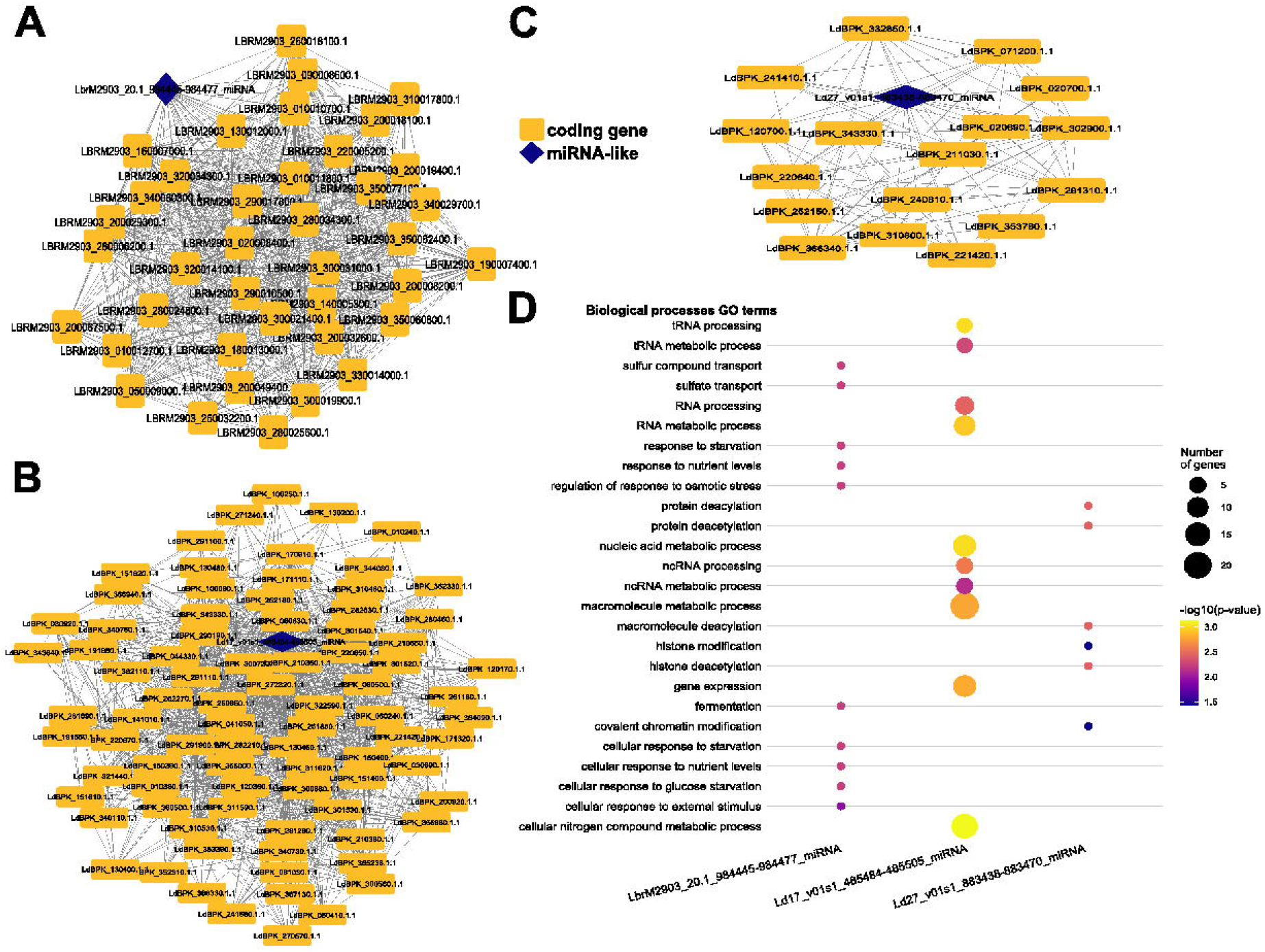
**A**. Sub-network of miRNA-like LbrM2903_20.1_984445-984477_miRNA (*L. braziliensis*). The grey triangle represents miRNA-like LbrM2903_20.1_984445-984477_miRNA. The grey circles represent the protein-coding genes that are co-expressed with it. B. Sub-network of miRNA-like Ld27_v01s1_883438-883470_miRNA (*L. donovan*i). C. Subnetwork of miRNA-like Ld17_v01s1_485484-485505_miRNA (*L. donovani*). **D**. Top 10 biological processes terms for functions assigned to these miRNAs-like.

## DISCUSSION

Here we have described for the first time a genome-wide prediction of non-coding RNAs in 16 species from the *Leishmania* genus. Our results on 26 different strains revealed expression patterns of coding and non-coding RNAs highly related to different parasite developmental stages. These findings may suggest a role in the regulation of morphological differentiation in both insect and mammal host stages, as occurs in other Trypanosomatidae (62). So far, different efforts have been performed in order to describe the repertory of distinct ncRNA classes in *Leishmania* parasites (44–46,48,49,51,52). However, different from these previous works, our computational approach did not focus on any particular ncRNA class, thus allowing an unbiased genome-wide identification of the ncRNAs repertoire. We followed a similar approach implemented before by our group to describe the sets of ncRNAs available in *Leishmania braziliensis* (51).

We combined sequence similarity searches and covariance model comparisons in order to identify and annotate a large number of ncRNAs in all studied genomes. Additionally, we used publicly available RNA-seq assays to obtain transcriptional evidence to validate these predictions. The results revealed a majority of siRNA-like with length varying between 20 to 25 nt, which coincides to the previously described for *L. braziliensis* (45). Additionally, as reported in *L. major*, most miRNA-like elements possess a length varying around 20 to 26 nt (46). Given our results shown heterogeneity of size and number of predicted ncRNA observed in all species, we identified a median size of 23 nt for all these putative ncRNAs. This finding contrasts greatly with the data presented by Ruy and colleagues (2019), who observed a median size of 281 nt in their search in *L. braziliensis*. Interestingly, when comparing our method with other approaches, we realized that their rationale was based on the identification of non-coding transcripts and then asked for an RNA class. Instead, we based our search on sequence alignment and probabilistic models and then verified our findings using RNA-seq data. Ruy colleagues (2019) also found more lncRNAs than we did with our approach and our results showed a larger proportion of small ncRNAs (52). This may suggest that they were identifying primary transcripts that could be precursors of small RNA classes identified in our analysis, which will require further analysis to be verified.

The conservation of ncRNA sequences has been observed in phylogenetically distant clades, such as between *Caenorhabditis elegans* and *Homo sapiens (63)*, even in an inter-kingdom level, such as the miR485 family (64), which is conserved between *Arabidopsis thaliana, C. elegans, Mus musculus*, and *H. sapiens*; as well as in more closely related species, such as *A*. and *A. lyrata* (65). This conservation has been firmly related with development, such as in nervous system and brain development, presenting orthologous within birds (*Gallus gallus*), marsupial (*Monodelphis domestica*), and eutherian mammals (*M. musculus*) (66,67), or in plant reproductive development (68); with diseases, such as neurodegenerative diseases or viral infections (69–71); or to stress responses, such as starvation, growth arrest and arsenic induced stress (72–76). Considering the accepted phylogenetic relationship between the *Leishmania* genus and their potential to infect different hosts, as well as the ability of this parasite to be associated with many clinical manifestations, we studied a possible relationship between sequence conservation of different ncRNAs classes and their role in parasite development, by comparing their expression profile in different developmental stages. The results we obtained here yields evidence of ncRNAs great conservation in *Leishmania* parasites among different species. These 436 core clusters represent more than 70% of all predicted ncRNAs, leaving a small proportion of non-conserved ncRNAs. Additionally, as expected, we observed that conservation increases as phylogenetic distance decreases (Supplementary Figure 1). Our analysis also identified an increase in ncRNA conservation in species that are canonically related to the same disease pathology or subgenus (Figure 2C), as previously reported for *Leishmania* parasites (47,52) and other trypanosomatids (77). Sequence conservation is also an indicator of selective pressure, as occurs with other species, and it can be associated with shared biological processes within related organisms (78). Previous studies on other pathogens showed that many ncRNA classes are related to different developmental steps (79,80). Thus, we observed that ncRNAs conservation may indicate an important role in the development and pathogenesis of *Leishmania* parasites.

The miRNA/RNAi pathway has been divided into a canonical and a non-canonical pathway (81–83). The canonical one requires the presence of Drosha, DGCR8, Dicer, Exportin 5 and Argonaute proteins (81–83). In contrast, there are several non-canonical miRNA biogenesis pathways described, and these are divided into Drosha-independent and Dicer-independent pathways (81–83). Since the RNAi mechanism was described in *Trypanosoma brucei* (84) and *Leishmania braziliensis* (85), little is known about the whole pathway in *Leishmania* parasites. A study conducted by Padmanabhan and cols. (2012) revealed the presence of AGO-like proteins in *Leishmania*, lacking the central domain PAZ, without activity in the biogenesis of ncRNAs (86). Although, RNAi mechanism was described 12 years ago in *L. braziliensis (85)*, only the AGO-1 family proteins were characterized and no other proteins of this biogenesis pathway were predicted or investigated. In this work, we made a computational approach to identify all the other proteins involved in the RNAi/miRNA biogenesis pathway. Interestingly, with our search approach based on functional domains and remote homology, we did not find evidence of the presence of Drosha, DGCR8 and TRBP proteins, but we did identify a putative exportin-5 (XPO5) gene in the genomes of *L. braziliensis, L. donovani* and *L. major*. Exportin-5 is essential for the canonical biogenesis RNAi pathway, and it is required for the nuclear exportation (87,88). These discoveries lead us to propose that a non-canonical pathway in which Drosha and DGCR8 do not participate could be used by *Leishmania* species. This is reaffirmed due to the characterization of small RNAs derived from both tRNAs and rRNAs, described in *L. donovani* and *L. braziliensis* (47). We further investigated the putative non-canonical RNAi/miRNA pathways present in *Leishmania* species. Here we identified a possible mechanism that involves m7G cap modification, which is a modification present in all transcripts of *Leishmania* species. This modification is performed by MTR1 in mRNA transcripts and MTTL1 in tRNAs. Interestingly, MTTL1 promotes biogenesis by 7-methylguanosine (m7G)-capped pre-miRNA in mammals (89), and several miRNAs, including 7-Let, a miRNA involved in development and tumor repression (90,91) which is known to be processed through this mechanism. However, the existence of this pathway has not been validated in *Leishmania* parasites and more work is needed for the characterization of the role of MTTL1 in miRNA processing in these parasites.

In order to understand the differences in gene expression along distinct developmental stages in *Leishmania* parasites, we selected three representative strains according to their disease type, subgenus and availability of RNA-seq datasets in public databases. In this sense, we compared the expression of both coding genes and ncRNAs from *L*. (*Viannia*) *braziliensis*, which causes cutaneous and mucocutaneous leishmaniasis; *L*. (*Leishmania*) *donovani*, which causes visceral leishmaniasis; and *L*. (*L*.) *major*, that causes cutaneous leishmaniasis. Our results show an expression of several proteins involved in host-parasite interaction in *L. braziliensis* amastigotes, such major surface protease GP63 (leishmanolysin), a protein that are involved in the survival of intracellular amastigotes (92). Also, this protein allows promastigotes to evade the complement-mediated lysis prior to its internalization by macrophages (93). The expression of this gene was also observed in metacyclic promastigotes in *L. braziliensis*. Additionally, we also identified Biopterin Transporter (BT1) overexpression in this stage, which product plays a key role in growth, infectivity and survival in the macrophages (94). In two computational studies, the potential of the GP63 protein from *L. major* and *L. donovani* as a vaccine target was demonstrated (95,96). Another study performed by Chowdhury and cols. (2019), designed two siRNAs and three miRNA that had *L. donovani* GP63 as their exclusive target, demonstrating that these molecules can be used to inhibit the expression of GP63 and act as one more therapeutic tool for tackling leishmaniasis (97). This could inspire more tests to be performed on GP63 and to observe its potential as a vaccine and therapeutic target.

Our study was based on the identification of miRNA-like gene functions associated with amastigote developmental stage, based on a coding/non-coding gene coexpression network. This method has been employed to determine the function of different classes of ncRNAs, such lncRNAs and miRNAs, in distinct etiological agents of infectious diseases, such as *Plasmodium falciparum* (61), *Toxoplasma gondii* (98) and *Schistosoma mansoni* (99). These works utilize different combinations of methods to determine the functional assignment. Based on this, we applied a topology and expression filter to predict the function of the top ten hub miRNA-like in amastigote developmental stage in *L. braziliensis* and *L. donovani*. Here, we identify several biological processes of interest, among which we can mention the histone modification in the top ten terms of *L. donovani* and the response to starvation and nutrient deficit in the top ten of *L. braziliensis*. Due to the annotated functions, we decided to increase the network distance to link genes to these ncRNAs, obtaining three miRNA-like; one for *L. braziliensis* and two for *L. donovani* that were associated with these predicted functions. Among the neighbors of these miRNAs in their network, we highlight TOR3 (target of rapamycin kinase kinase 3), a protein that belongs to a family implicated in nutrient stress (100). However in *Leishmania* TOR3 protein is essential for survival and infection in macrophages and it is known to be implicated in stress response (101–104). The relevance of this protein in the infection and survival and their co-expression with the miRNA-like gene LbrM2903_20.1_984445-984477_miRNA, indicate that the expression of this RNA could play a crucial role in *L. braziliensis* survival.

Histone deacetylases (HDACs) are a family of proteins that act in transcriptional repression by histone deacetylation (105). In trypanosomatids HDACs have been proposed as a potential therapeutic target (106) and its inhibition shows a reduction in the infection and replication of intracellular amastigotes (107). Here, we documented the co-expression of Ld27_v01s1_883438-883470_miRNA with HDAC in *L. donovani* amastigotes, which may be indicating a possible relation of this miRNA-like with the replication and infection establishment. Some studies have shown that histone proteins are evolutionarily diversified in trypanosomatids in comparison with histones from other eukaryotes (34,108,109). Several previous works demonstrated that histone, specially H2B and H4, show antigenicity activity and were recognized by antibodies (110), apart from being upregulated when treated with sodium antimony gluconate (SAG) (111). The presence of anti-H2B antibodies has been found in symptomatic visceral and cutaneous leishmaniasis patients (112,113), corroborating that histones are needed for the correct development and survival of the parasite. Additionally, this indicates that the Ld17_v01s1_485484-485505_miRNA (form *L. donovani*), which co-expresses with histone H2B, may play an important role in the regulation of parasite’s development and survival in hosts by regulating histones and other proteins. Thus, we propose histone H2B and the co-expressed miRNA-like genes as possible markers of drug susceptibility and survival.

In summary, our work is the first to identify novel ncRNAs in 26 genome isolates from 16 *Leishmania* species depicted from its pan-RNAome. We revealed a new set of non-coding genes that are involved in different developmental stages in the parasite life cycle and may play an important role in infection and survival. Furthermore, we obtained the co-expression profile of coding and non-coding genes functional insights for the potential role of several miRNAs-like through guilt-by-association inferences. Importantly, our results provide novel evidence of possible mechanisms underlying co-regulation between coding-miRNA-like interaction in the regulation histone deacetylation, opening the way for further research on the role of these ncRNAs and their possible relationship with parasite survival.

## METHODS

### Databases and datasets

All genome sequences of 26 strains from 16 different species of *Leishmania* spp. were downloaded from NCBI (114) FTP site and TriTrypDB release 35 (115), in October 2019. Publicly available RNA sequencing (RNA-seq) libraries were downloaded from SRA NCBI (116) in October 2019. Samples from three *Leishmania* species at different developmental stages were collected: *L. braziliensis* M2903 (MHOM/BR/1975/M2903) study accession PRJNA494068 was published by Ruy and cols. (2019) (52) and containing samples for amastigote, procyclic and metacyclic developmental stages. *L. donovani* BPK282A1 (MHOM/NP/2002/BPK 282) study accession PRJEB15610 was published by Dumentz and cols. (2017) (19), which contains information on amastigote, undifferentiated promastigote; and *L. major* Friedlin (MHOM/IL/1981/Friedlin) study accession PRJNA252769 was published by Fernandes and cols. (2016) (56) and contains samples for metacyclic and procyclic.

### Predicting the repertoire of ncRNAs in *Leishmania* spp. genomes

Predictions of ncRNAs in *Leishmania* spp. genomes were carried out using two different schemes. The first approach is based on sequence homology detection, using the set of non-redundant sequences available at the NR2 database (54). For this, we employed an in-house developed script to perform the search. This script takes as input a FASTA file with all sequences from NR2 and a genome index made with Bowtie2 (117) for each genome, generating as output a BED file for each chromosome with all mapped ncRNAs. The other ncRNA prediction scheme was based on covariance models search, as implemented in StructRNAFinder (118), using Rfam 14.1 (55) known RNA families. An e-value cut-off of 0.001 was applied for cmsearch and a score of 10 was used to identify and annotate RNA families with StructRNAfinder, as previously applied for *L. braziliensis* (51). The two BED files obtained from both sequence homology and covariance models approaches were merged, in order to eliminate redundancies, using the BEDtools merge (119). Intragenic ncRNAs were eliminated using intersectBED (119). The final set of ncRNA FASTA sequences was obtained using the BEDtools getfasta (119). GTF files for all predicted ncRNAs were builded using UCSC tools (https://github.com/ucscGenomeBrowser/kent). The final RNA class annotation for each predicted ncRNA was obtained by using the consensus made between the classes that were mapped by both searches.

### Transcriptional evidence of predicted ncRNAs

Once the putative ncRNAs were identified, their expressions were used to validate their existence. We selected *Leishmania braziliensis* MHOM/BR/1975/M2903, *Leishmania donovani* BPK282A1 (MHOM/NP/2002/BPK282A1) and *Leishmania major* Friedlin (MHOM/IL/1981/Friedlin) for subsequent analyzes, according to previously described parameters (47). Next, we employed an automated in-house pipeline for RNA-seq mapping and read counts measuring (120), with the following modifications. FastQC v0.11.8 (https://www.bioinformatics.babraham.ac.uk/projects/fastqc/) and Fastp (121) were used to evaluate and filter out low quality reads, considering a Phred cut-off value of Q = 30. Then, we mapped the remaining high quality reads using Bowtie2 (117) to each one of the three selected genomes. Expression read count values for all predicted ncRNAs were calculated using HTSeq-count 0.7.2 (122) and normalized using CPM from edgeR (123), for this the ncRNA gtf file was then added to the public protein coding gtf file of each specie. A CPM cut-off value of 0.5 was employed to determine if a particular ncRNA prediction was deemed to exist in each genome.

### Comparative analysis of ncRNAs

The ncRNA repertoire in *Leishmania* spp. was analyzed to determine their conservation within all studied genomes. First, we recovered the FASTA sequence for all ncRNA and clustered using CD-HIT version 4.8.1 (57), considering 80% of sequence similarity and coverage of both largest and shortest sequences to classify an ncRNA as a cluster member. Then, a binary presence-absence matrix was constructed based on each cluster of predicted ncRNAs. The set of ncRNA clusters shared within all strains was defined as the “core-RNAome”, while the sum of the “dispensable RNAome” (strain specific (unique) and accessory ncRNA clusters) and the core RNAome, was defined as the “pan-RNAome”.

To visualize the species clustering according to the ncRNA family distribution, we applied a hierarchical clustering with Euclidean distance using the presence/absence matrix. Next, we compared the generated cladogram with a phylogenetic tree obtained from a set of conserved orthologous proteins. For that, core-genome phylogenetic trees were built using 613 conserved orthologous proteins from the core-genome of 26 *Leishmania* representatives. We employed a bidirectional best-hit algorithm for orthologs clustering using ProteinOrtho v5.11 (124) with default parameters. Next, conserved orthologous multiprotein family sequences were aligned using MAFFT version v7.453 (125), considering the L-INS-I iterative refinement method. The alignments were masked to remove unreliable aligned regions with GBLOCKS version 0.91b (126). Maximum likelihood trees were prepared for concatenated alignments through IQ-TREE version 1.6.12 (127), using 1000 replicates as bootstrap with best-suited substitution model. The final tree was visualized using Figtree (http://tree.bio.ed.ac.uk/software/figtree/).

### Computational characterization of RNAi/miRNA pathway

With the aim to characterize the RNAi/miRNA pathway in Leishmania genomes, we applied a computational search based on two schemes. First all protein components for both canonical and non-canonical RNAi/miRNA biogenesis pathways previously described by Miyoshi and cols. (2010) (82), Abdelfattah and cols. (2014) (83) and Sand (2014) (128) was selected, then the Pfam ids of all domains of these proteins were obtained and their Hidden Markov Models (HMMs) were downloaded from Pfam v33.1 (129). After that, we scan all predicted proteins in *Leishmania* genomes against downloaded HMMs using hmmscan from HMMER v3.3 (http://hmmer.org/) (130). The cutoff value employed for domain annotation was 1E-04 E-value used by Hu and cols. (2019) to determine DNA binding domains in transcription factors (131).

Next, to determine the presence of AGO-like proteins in *L. donovani* and *L. major* species, we performed a local alignment with BLAST v2.9.0 (132) using as a database all AGO-like proteins reported in TritrypDB (115) and AGO proteins obtained from the previous step. Cutoff values employed in this search were 50% of identity, 77% of query coverage and e-values <1e-10.

### Co-expression network analysis

Weighted gene co-expression network analyses were performed to identify modules related to the developmental stages using WGCNA tool version 1.69 (60). Only genes with expression greater than 0.5 counts per million (CPM) in at least 90% of the libraries in one or more developmental stages were considered. Expression levels then were transformed into log space using log2 transform from EdgeR (123). The adjacency matrix was built using the power adjacency function for unsigned networks and was applied with the soft-thresholding beta parameter equal to 14 for *L. braziliensis* and *L. donovani*, and a beta = 22 for *L. major*; which resulted in a scale-free topology model fit index (R2 = 0.85). The adjacency matrix was then converted to the Topological Overlap Matrix (TOM) and the dissimilarity TOM (1 − TOM) was calculated to identify hierarchical clustering nodes and modules (99,133). Module-developmental stage correlation was calculated using Pearson coefficient between the expression levels of the genes belonging to each module along the developmental stages (99). Next, to analyze the correlation of each individual gene presenting high correlation to different developmental stages, we calculate gene significance (GS) and module membership (MM) mesures. By assigning a gene *X* as highly related to developmental stage *Z*, if an X(Z)= GS > 0.70, *p* < 0.05 and MM > |0.5|, *p* < 0.05.

### Differential expression analysis among developmental stages in *Leishmania* spp

The comparison of differential expression values between all developmental stages of *Leishmania* parasites was generated using DESeq2 v1.34.0(98). The differential screening parameter used as cutoff to detect a differentially expressed gene (DEG) was a *p value* < 0.05, false discovery rate (FDR) < 0.001 and |log2FC| > 0.75. A set of candidate genes that are more related to some of the developmental stages in *Leishmania* parasites were identified by overlapping the DEGs and the genes in the key modules.

### Functional associations of ncRNA in *Leishmania* parasites

To obtain functional associations for each ncRNA we employed a modified version of the method described by Liao and cols (61). First, we obtained the Gene Ontology (GO) annotations GAF files from TritrypDB (115), which contains the GO term annotations for each gene, including biological process (BP), cellular component (CC), and molecular function (MF), Then, we calculated the *p*-value of each GO term, of the first-level coding neighbors for each ncRNA using a hypergeometric test with GOseq v1.48.0 (134). The *p*-values were adjusted using the FDR method and *p* < 0.05 was used as cut off. To facilitate the analysis, interpretation of GO terms were fused into their level-3 ancestors in the hierarchy.

## ACKNOWLEDGMENTS

We acknowledge the kind support received from Dr. Evandro Ferrada, Dr Fernando Alfaro and Dr Roberto Mercado. Their patience and useful discussions were valuable for the improvement of our work. Also, we would like to thank the members of both NBL and LIB laboratories for their constructive criticism and advice along the development of this manuscript.We thank Dr. Adriana Ludwig - Instituto Carlos Chagas, Fundação Oswaldo Cruz, Curitiba, PR, Brasil, for her discussion and advice and for sharing her background and thoughts on ncRNA in trypanosomatids. PhD scholarship from Universidad Mayor to J.E.M.H. RMN research is supported by Fundação de Amparo à Pesquisa do Estado de Minas Gerais – Fapemig, grant PPM-00699-18. RMN is CNPq - Conselho Nacional de Desenvolvimento Científico e Tecnológico - research fellow (grant code 312965/2020-6). AJMM and VMC research is funded by Agencia Nacional de Investigación y Desarrollo de Chile,grants FONDECYT Regular 1181089 and 1211731. Powered@NLHPC: this research was partially supported by the supercomputing infrastructure of the NLHPC (ECM-02); and by the computing infrastructure of the Centro de Genómica y Bioinformática, Universidad Mayor.

## SUPPORTING INFORMATION CAPTIONS

**Supplementary Table 1**. non-coding RNA prediction and GC content compared vs genome sequence

**Supplementary Table 2**. Number of all classes of non coding RNAs found in genomes of *Leishmania* spp.

**Supplementary Table 3**. Number of coding genes and ncRNAs per module in different *Leishmania* parasites

**Supplementary Figure 1. Phylogenetic analysis of *Leishmania* genus and comparative with presence/absence clustering of core RNAome**. Phylogenetic tree based on 613 core genes was compared against a dendrogram derived from Jaccard Coefficient Cluster analysis based on a similarity matrix obtained from pan-RNAome presence/absence binary matrix.

**Supplementary Figure 2. Differentially expressed coding and non-coding genes through developmental stages in *L. donovani*. A**. Pairwise comparison of gene expression in Amastigote and Promastigote developmental stages. The number on top of each bar represents the number of all overexpressed genes in each stage comparison. **B**. Top 10 biological processes terms according to *p*-value for each developmental stage.

**Supplementary Figure 3. Differentially expressed coding and non-coding genes through developmental stages in *L. major*. A**. Pairwise comparison of gene expression in Metacyclic and Procyclic developmental stages. The number on top of each bar represents the number of all overexpressed genes in each stage comparison. **B**. Top 10 biological processes terms according to p-value for each developmental stage.

**Supplementary Figure 4. Co-expression module-Developmental stages associations**. Each row corresponds to a module, column to a developmental stage. Each cell contains the corresponding correlation and p-value (parenthesis). The table is color-coded by correlation according to the color legend. **A**. *L. donovani*, module M11 (R^2^= 0.95 and *p* < 0.001) is narrowly associated with promastigotes and M5 (R^2^= 0.95 and *p* = 0.0011) with amastigotes **B**. *L. major*, module M2 (R^2^= 0.81 and *p* < 0.001) is correlated to procyclic and M9 (R^2^= 0.65 and *p* = 0.0023) to metacyclic.

**Supplementary File 1**. Differential Expressed Genes through developmental stages of Leishmania parasites

**Supplementary File 2**. GO enrichment of DEGs in *Leishmania* parasites **Supplementary File 3**. GO functional assignment to top 10 miRNA-like associated with Amastigote in *L. braziliensis* and *L. donovani*.

