## supplementary tables for "Comparative and systems analyses of *Leishmania* spp. non-coding RNAs through developmental stages"

| **Supplementary Table 1.** non-coding RNA prediction and GC content compared vs genome sequence | | | | | | | |
| --- | --- | --- | --- | --- | --- | --- | --- |
| Specie | Accession | Subgenus | Disease type | # Scaffolds | # of ncRNAs | GC% genomic | GC% ncRNAs |
| *Leishmania aethiopica* L147 (LaetL147) | GCA_000444285.2 | Leishmania | CL | 160 | 12560 | 58.66 | 57.8 |
| *Leishmania amazonensis* M2269 (LamaM2269) |  | Leishmania | CL | 2627 | 7094 | 58.4 | 57.54 |
| *Leishmania arabica* LEM1108 (LaraLEM118) | GCA_000410695.2 | Leishmania | ND | 168 | 14207 | 58.04 | 58.86 |
| *Leishmania braziliensis* M2903 (LbraM2903) | GCA_000340355.2 | Viannia | MCL | 745 | 10511 | 57.15 | 58.29 |
| *Leishmania braziliensis* M2904 (LbraM2904) | GCA_000002845.2 | Viannia | MCL | 139 | 10325 | 59.24 | 57.97 |
| *Leishmania donovani* AG83 (LdonAG83) | GCA_001989975.1 | Leishmania | VL | 36 | 6646 | 53 | 58.31 |
| *Leishmania donovani* BHU1220 (LdonBHU1220) | GCA_000470725.1 | Leishmania | VL | 36 | 9205 | 57.78 | 57.48 |
| *Leishmania donovani* BPK282A1 (LdonBPK282A1) | GCA_000227135.2 | Leishmania | VL | 36 | 9335 | 57.83 | 57.87 |
| *Leishmania donovani* Pasteur (LdonPasteur) | GCA_002243465.1 | Leishmania | VL | 37 | 10241 | 59 | 58.9 |
| *Leishmania enrietti* LEM3045 (LenrLEM3045) | GCA_000410755.2 | Mundinia | ND | 495 | 9600 | 58.87 | 54.84 |
| *Leishmania gerbilli* LEM452 (LgerLEM452) | GCA_000443025.1 | Leishmania | CL | 142 | 13981 | 57.91 | 59.27 |
| *Leishmania infantum* JPCM5 (LinfJPCM5) | GCA_000002875.2 | Leishmania | VL | 70 | 5149 | 61.14 | 59.5 |
| *Leishmania infantum* TR01 (LinfTR01) | GCA_003020905.1 | Leishmania | VL | 36 | 9068 | 60.15 | 58.22 |
| *Leishmania major* Friedlin (LmajFriedlin) | GCA_000002725.2 | Leishmania | CL | 36 | 15238 | 60.31 | 55.09 |
| *Leishmania major* LV39c5 (LmajLV39c5) | GCA_000331345.1 | Leishmania | CL | 809 | 14403 | 57.13 | 55.97 |
| *Leishmania major* SD75 (LmajSD75) | GCA_000250755.2 | Leishmania | CL | 36 | 15868 | 59.97 | 54.53 |
| *Leishmania mexicana* U11003 (LmexU1103) | GCA_000234665.4.1 | Leishmania | CL | 36 | 10321 | 61.29 | 59.8 |
| *Leishmania panamensis* L13 (LpanL13) | GCA_000755165.1 | Viannia | CL | 856 | 9744 | 54.41 | 60 |
| *Leishmania panamensis* PSC1 (LpanPSC1) | GCA_000340495.1 | Viannia | CL | 35 | 9245 | 56.91 | 59.09 |
| *Leishmania peruviana* LEM1537 (LperLEM1537) | GCA_001403675.1 | Viannia | CL | 37 | 10040 | 39.32 | 56.35 |
| *Leishmania peruviana* PAB4377 (LperPAB4377) | GCA_001403695.1 | Viannia | CL | 37 | 10157 | 52.29 | 57.09 |
| *Leishmania* sp LD974 (LspLD974) | GCA_000981925.2 | Unknown | PKDL | 1100 | 5340 | 54.71 | 47.78 |
| *Leishmania* sp LEM2494 (LspLEM2494) | GCA_000409445.2 | Mundinia | CL | 251 | 7753 | 59.39 | 58.2 |
| *Leishmania tarentolae* ParTarII (LtarParTarII) | GCA_009731335.1 | Sauroleishmania | ND | 1351 | 9484 | 56.27 | 48.63 |
| *Leishmania tropica* L590 (LtroL590) | GCA_000410715.1 | Leishmania | CL | 160 | 12857 | 57.1 | 58.14 |
| *Leishmania turanica* LEM423 (LturLEM423) | GCA_000441995.1 | Leishmania | CL | 219 | 17665 | 57.93 | 59.38 |
| CL = cutaneous leishmaniasis  MCL = mucocutaneous leishmaniasis  VL = visceral leishmaniasis  PKDL = Post-kala-azar dermal leishmaniasis (VL)  ND = non reported in humans | | | | | | | |

| **Supplementary Table 2**. Number of all class of non coding RNAs found in genomes of *Leishmania* spp | | | | | | | | | | | | | | | | | | | |
| --- | --- | --- | --- | --- | --- | --- | --- | --- | --- | --- | --- | --- | --- | --- | --- | --- | --- | --- | --- |
| Specie | ncRNA Class | | | | | | | | | | | | | | | | | | |
|  | Cis-reg | gRNA | HACA-box | IRES | lncRNA | miRNA | piRNA | ribozyme | rRNA | RUF21 | scaRNA | siRNA | snoRNA | snRNA | sRNA | SRPRNA | ta-siRNA | tRNA | Unclassified |
| LaetL147 | 2 | 1 | 1 | 0 | 3 | 1847 | 896 | 0 | 364 | 0 | 0 | 121 | 19 | 3 | 1221 | 0 | 0 | 82 | 8000 |
| LamaM2269 | 9 | 1 | 1 | 0 | 7 | 1364 | 485 | 0 | 137 | 0 | 0 | 123 | 11 | 5 | 938 | 0 | 0 | 50 | 3963 |
| LaraLEM1108 | 1 | 1 | 1 | 0 | 1 | 1935 | 988 | 0 | 412 | 0 | 0 | 180 | 8 | 3 | 1451 | 0 | 0 | 81 | 9145 |
| LbraM2903 | 14 | 1 | 0 | 0 | 6 | 1889 | 776 | 0 | 531 | 0 | 0 | 133 | 38 | 4 | 1059 | 0 | 0 | 76 | 5984 |
| LbraM2904 | 15 | 0 | 0 | 0 | 2 | 1832 | 678 | 0 | 540 | 0 | 0 | 136 | 38 | 2 | 967 | 0 | 0 | 99 | 6016 |
| LdonAG83 | 2 | 0 | 0 | 0 | 0 | 1126 | 382 | 0 | 142 | 0 | 0 | 92 | 7 | 1 | 707 | 0 | 0 | 32 | 4155 |
| LdonBHU1220 | 3 | 0 | 0 | 0 | 0 | 1552 | 523 | 0 | 195 | 0 | 0 | 125 | 16 | 2 | 950 | 0 | 0 | 48 | 5791 |
| LdonBPK282A1 | 3 | 0 | 0 | 0 | 0 | 1561 | 551 | 0 | 198 | 0 | 0 | 125 | 16 | 2 | 953 | 0 | 0 | 54 | 5872 |
| LdonPasteur | 5 | 1 | 0 | 0 | 1 | 1703 | 602 | 0 | 210 | 0 | 0 | 114 | 42 | 2 | 1051 | 0 | 0 | 75 | 6435 |
| LenrLEM3045 | 2 | 1 | 0 | 0 | 1 | 1138 | 690 | 0 | 479 | 0 | 0 | 103 | 4 | 7 | 757 | 0 | 0 | 42 | 6376 |
| LgerLEM452 | 2 | 1 | 0 | 0 | 0 | 2201 | 994 | 0 | 421 | 0 | 0 | 168 | 8 | 3 | 1438 | 0 | 0 | 64 | 8681 |
| LinfJPCM5 | 4 | 0 | 0 | 1 | 0 | 930 | 302 | 0 | 119 | 0 | 0 | 87 | 19 | 0 | 526 | 0 | 0 | 30 | 3131 |
| LinfTR01 | 3 | 0 | 0 | 0 | 0 | 1621 | 538 | 0 | 170 | 0 | 0 | 141 | 21 | 2 | 919 | 0 | 0 | 61 | 5592 |
| LmajFriedlin | 5 | 0 | 0 | 0 | 0 | 1963 | 1160 | 0 | 806 | 0 | 0 | 159 | 22 | 2 | 1298 | 0 | 0 | 69 | 9754 |
| LmajLV39c5 | 2 | 1 | 0 | 0 | 1 | 1849 | 1070 | 0 | 770 | 18 | 0 | 164 | 19 | 6 | 1202 | 0 | 0 | 63 | 9238 |
| LmajSD75 | 2 | 0 | 0 | 0 | 0 | 1760 | 1121 | 0 | 790 | 0 | 0 | 1447 | 14 | 2 | 1244 | 0 | 0 | 61 | 9427 |
| LmexU1103 | 8 | 1 | 1 | 0 | 0 | 1813 | 648 | 0 | 231 | 0 | 0 | 139 | 24 | 5 | 1222 | 1 | 0 | 83 | 6145 |
| LpanL13 | 19 | 1 | 0 | 0 | 8 | 1880 | 888 | 0 | 513 | 0 | 0 | 115 | 24 | 2 | 1048 | 0 | 0 | 70 | 5176 |
| LpanPSC1 | 14 | 0 | 0 | 0 | 6 | 1805 | 623 | 0 | 529 | 0 | 0 | 117 | 24 | 2 | 1008 | 0 | 0 | 76 | 5041 |
| LperLEM1537 | 13 | 0 | 0 | 2 | 3 | 1736 | 800 | 0 | 529 | 0 | 0 | 128 | 38 | 2 | 978 | 0 | 0 | 51 | 5760 |
| LperPAB4377 | 13 | 0 | 0 | 2 | 2 | 1807 | 724 | 0 | 543 | 0 | 0 | 120 | 36 | 3 | 992 | 1 | 0 | 73 | 5841 |
| LspLD974 | 1 | 0 | 0 | 0 | 1 | 644 | 355 | 2 | 99 | 0 | 1 | 113 | 13 | 31 | 700 | 1 | 0 | 42 | 3337 |
| LspLEM2494 | 3 | 0 | 0 | 0 | 0 | 1991 | 468 | 0 | 184 | 0 | 0 | 175 | 5 | 9 | 900 | 0 | 0 | 57 | 3961 |
| LtarParTarII | 3 | 79 | 0 | 0 | 2 | 1317 | 491 | 0 | 230 | 0 | 0 | 97 | 5 | 4 | 933 | 0 | 1 | 61 | 6261 |
| LtroL590 | 2 | 1 | 1 | 0 | 1 | 1661 | 737 | 0 | 432 | 0 | 0 | 114 | 15 | 2 | 1194 | 0 | 0 | 67 | 8630 |
| LturLEM423 | 1 | 0 | 0 | 0 | 0 | 2371 | 1287 | 0 | 808 | 0 | 0 | 177 | 19 | 2 | 1665 | 0 | 0 | 78 | 11257 |

| **Supplementary Table 3**. Number of coding genes and ncRNAs per module in different *Leishmania* parasites | | | | | | | | | |
| --- | --- | --- | --- | --- | --- | --- | --- | --- | --- |
| **Module** | ***L. braziliensis*** | | | ***L. donovani*** | | | ***L. major*** | | |
|  | **Total** | **coding** | **ncRNA** | **Total** | **coding** | **ncRNA** | **Total** | **coding** | **ncRNA** |
| **M1** | 749 | 540 | 209 | 1999 | 1066 | 933 | 107 | 4 | 103 |
| **M2** | 303 | 253 | 50 | 43 | 25 | 18 | 221 | 143 | 78 |
| **M3** | 172 | 138 | 34 | 220 | 119 | 101 | 5641 | 2540 | 3101 |
| **M4** | 124 | 106 | 18 | 513 | 216 | 297 | 190 | 116 | 74 |
| **M5** | 59 | 45 | 14 | 3398 | 2158 | 1240 | 102 | 88 | 14 |
| **M6** | 714 | 535 | 179 | 122 | 41 | 81 | 1458 | 589 | 1458 |
| **M7** | 427 | 300 | 127 | 191 | 112 | 79 | 712 | 237 | 475 |
| **M8** | 945 | 630 | 315 | 80 | 42 | 38 | 165 | 132 | 33 |
| **M9** | 838 | 544 | 294 | 1056 | 659 | 397 | 3439 | 1077 | 2362 |
| **M10** | 107 | 103 | 4 | 72 | 27 | 45 | 2600 | 1495 | 1105 |
| **M11** | 109 | 94 | 15 | 4896 | 2446 | 2450 | 74 | 72 | 2 |
| **M12** | 101 | 98 | 3 | 119 | 75 | 44 | 277 | 230 | 47 |
| **M13** | 55 | 55 | 0 | 139 | 73 | 66 | 336 | 16 | 320 |
| **M14** | 1130 | 867 | 263 | 297 | 130 | 167 | 782 | 192 | 590 |
| **M15** | 277 | 221 | 56 | 533 | 348 | 185 | 241 | 139 | 102 |
| **M16** | 204 | 192 | 12 | 105 | 30 | 75 | 130 | 37 | 93 |
| **M17** | 485 | 391 | 94 | 1298 | 431 | 867 | - | - | - |
| **M18** | 378 | 262 | 116 | - | - | - | - | - | - |
| **M19** | 1855 | 1423 | 432 | - | - | - | - | - | - |
| **M20** | 2539 | 1783 | 756 | - | - | - | - | - | - |
| **M0** | 1662 | 328 | 1334 | 1 | 0 | 1 | 2931 | 1008 | 1923 |
