## Supplementary figures and images for "Comparative and systems analyses of *Leishmania* spp. non-coding RNAs through developmental stages"

### Supplementary Figure 1

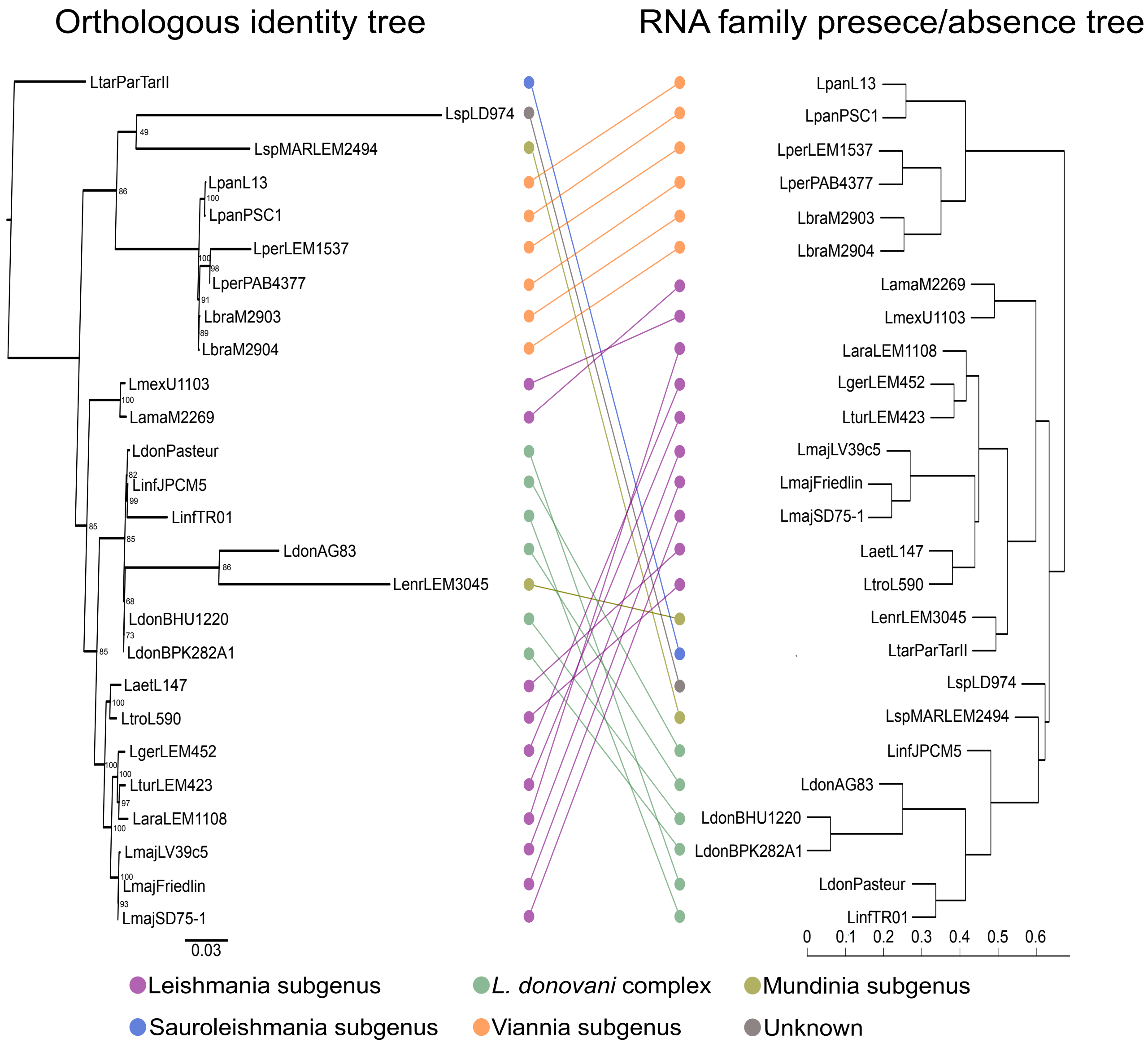

### Supplementary Figure 2

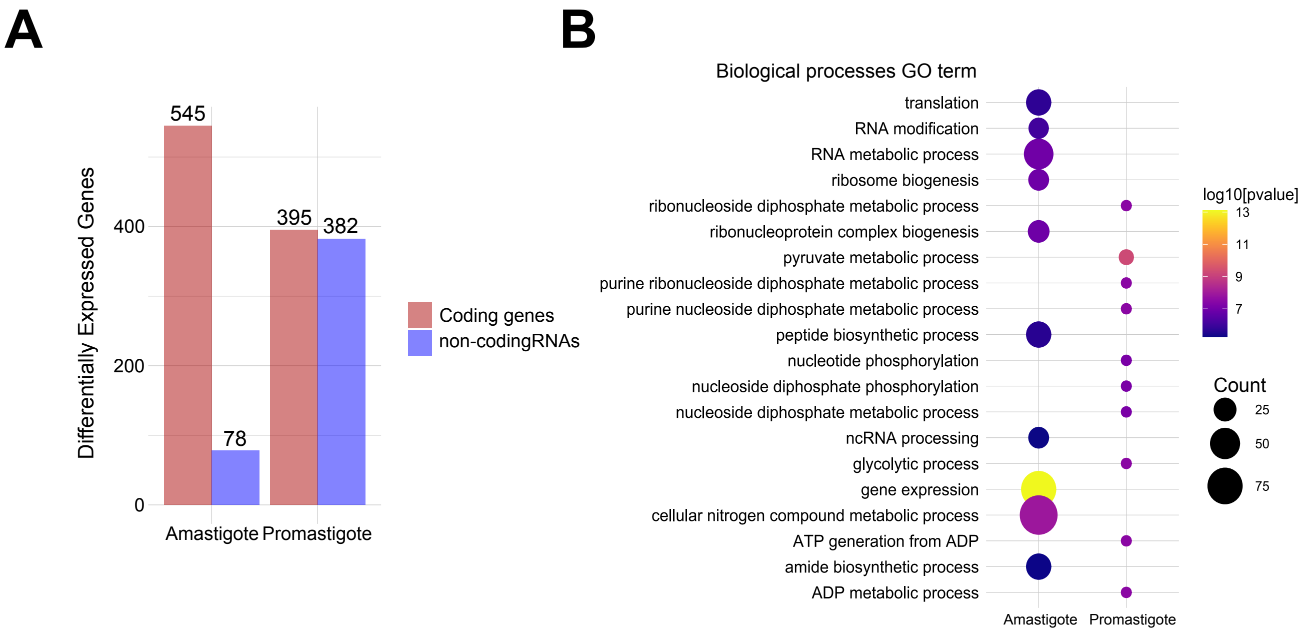

### Supplementary Figure 3

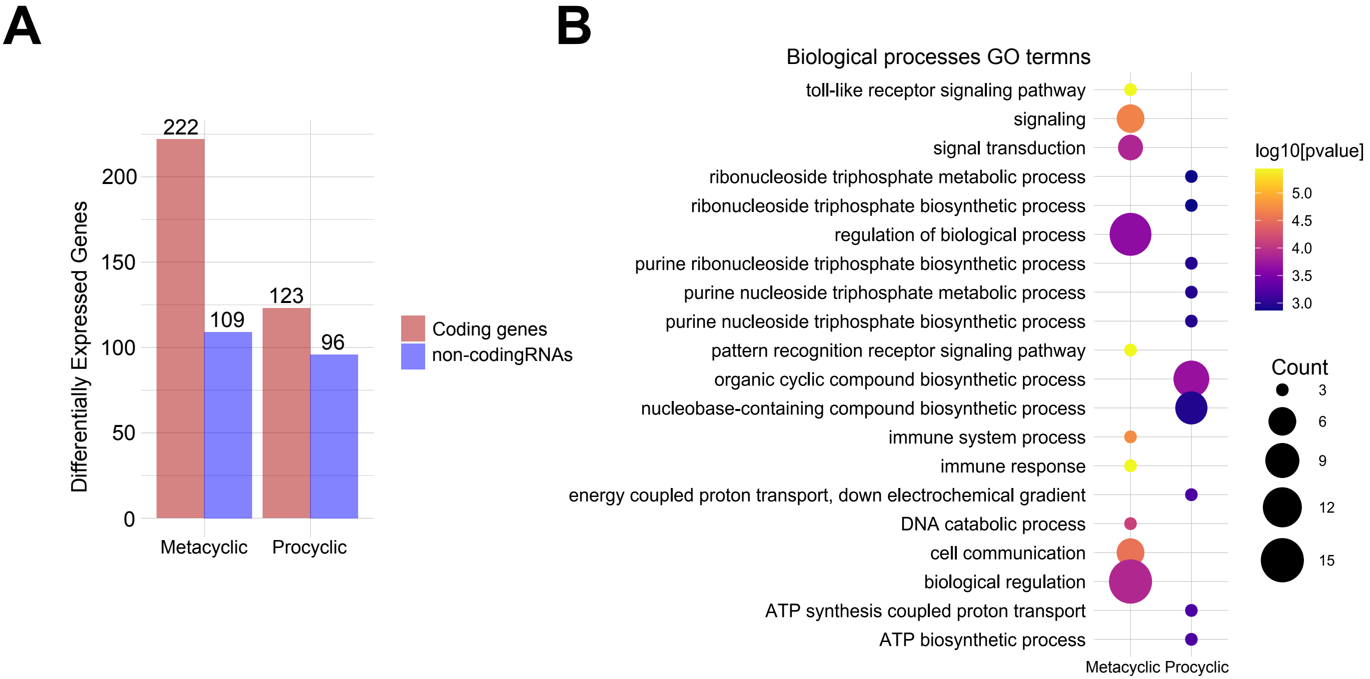

### Supplementary Figure 4

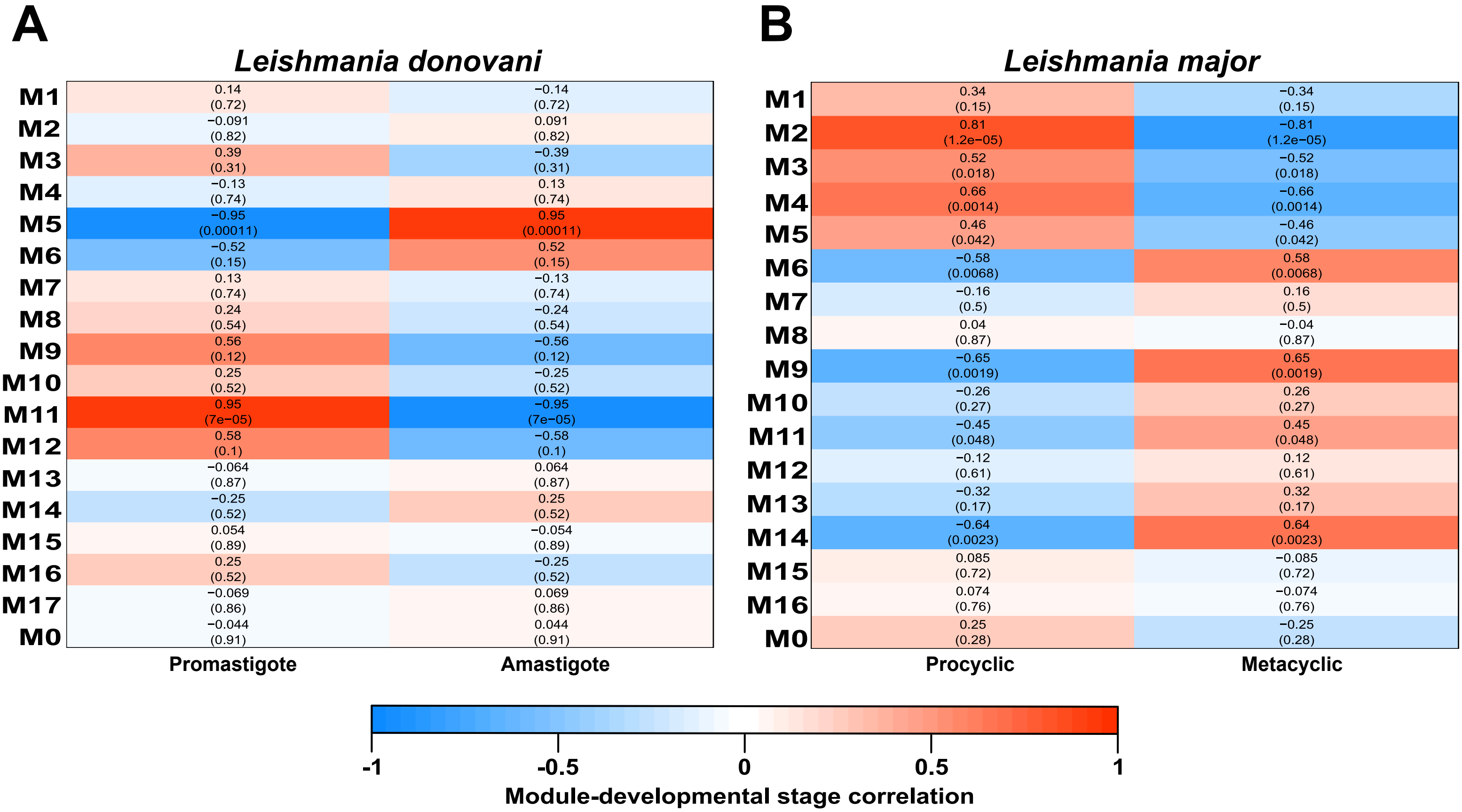
